# The truncate soft-shell clam, *Mya truncata*, as a biomonitor of municipal wastewater exposure and historical anthropogenic impacts in the Canadian Arctic

**DOI:** 10.1101/2021.03.29.437602

**Authors:** Christina M Schaefer, David Deslauriers, Ken M Jeffries

## Abstract

Municipal wastewater is a large source of pollution to Canadian waters, yet its effects on Arctic marine ecosystems remains relatively unknown. We characterized the impacts of municipal wastewater from a growing northern community, Iqaluit, Nunavut on the Arctic truncate soft-shell clam, *Mya truncata*. Clams were sampled from six locations that varied in proximity to the wastewater treatment plant and shell biogeochemical analysis revealed that clams nearest the wastewater treatment plant had slower growth rates, lower carbon and oxygen stable isotope ratios, and elevated concentrations of copper and lead. A parallel analysis on mRNA expression profiles characterized *M. truncata*’s physiological response to wastewater effluent. Clams nearest the wastewater treatment plant had significantly lower mRNA expression of genes associated with metabolism, antioxidants, molecular chaperones, and phase I and II detoxification, but had heightened mRNA expression in genes coding for enzymes that bind and remove contaminants. These results demonstrated a biological response to Iqaluit’s wastewater effluent and highlight *M. truncata*’s potential to act as a biomonitor of municipal wastewater along Canadian Arctic coastlines.

## Introduction

The discharge of municipal wastewater into aquatic ecosystems is a global practice and despite increasingly strict governmental guidelines, wastewater effluent contains a myriad of substances that may impact aquatic organisms (Gunnarsdóttir et al., 2013). Wastewater management is even more challenging in the Arctic as the harsh environment dictates the design, accessibility, construction, and operation of wastewater treatment plants (WWTP) (Daley et al., 2015). Local wastewater contamination in the Arctic has received little attention relative to long-range contaminants, but recent studies have shown that low water temperatures can reduce the breakdown of contaminants and deteriorate the quality of an already fragile coastal environment (Gunnarsdóttir et al., 2013). Wastewater effluent often consists of a high input of organic matter, detergents, pharmaceuticals, personal care products, polycyclic aromatic hydrocarbons (PAHs), surfactants, and trace metals; which could have strong influences on primary production, salinity, water temperature, and the overall structure of the coastal Arctic marine ecosystem (Falfushynska et al., 2014; Medeiros et al., 2008). Conventional monitoring of wastewater impacts on aquatic systems remains inadequate in the Arctic and this necessitates the use of environmental proxies like biomonitors to quantify the bioavailability of contaminants and assess the significance of contaminant exposure within the aquatic environment (Hatje, 2016; Phillips and Segar, 1986).

The Arctic truncate soft-shell clam, *Mya truncata*, has a wide northern distribution range and is long-lived, sessile, filter-feeding, and typically abundant in northern benthic communities (Sleight et al., 2018, 2016). These characteristics highlight the potential to establish this clam species as a biomonitor for investigating environmental variability in remote Arctic ecosystems. Moreover, this clam’s capability for contaminant accumulation is of great concern as these bivalves are recreationally harvested and serve as a culturally important natural food for northern Indigenous peoples (AMAP, 1998; Wakegijig et al., 2013).

The shells of the truncate soft-shell clam, and bivalves in general, retain an ontogenetic record of growth in the form of concentric growth lines that form in equal duration in undisturbed conditions (Schöne and Surge, 2012). As these bivalves grow, their shells increase in size and thickness by means of calcium carbonate deposition onto the organic matrix, a process termed biomineralization (Schöne and Surge, 2012). In addition to the building blocks required for biomineralization (i.e., calcium, bicarbonate, organic molecules), trace elements and stable isotopes present in the ambient water at the time of growth are also incorporated into the shell (Gillikin et al., 2005). As a result, the long lifespan of these clams provides a medium to track historical changes in environmental conditions through growth patterns, stable isotopes, and trace element signatures in the shell. This information is beneficial in remote ecological systems where historical records are not readily available and can be used to interpret spatial and temporal temperature, salinity, ocean productivity, and contamination patterns during the periods of growth (Jones and Quitmyer, 1996).

Molecular biomarkers (i.e., messenger RNA (mRNA) transcripts) can provide sensitive expression profiles to assess the shorter-term physiological impacts of pollutants from wastewater effluent on organisms (Li et al., 2013). Heat shock proteins (HSPs) are a highly conserved family of molecular chaperones, typically recruited as first responders to cellular stress (Fabbri et al., 2008). These HSP mRNA transcripts are induced by a variety of stimuli within wastewater effluent, rendering them indicators of the general health condition for the organism (Fabbri et al., 2008). Wastewater contaminants have also been shown to impact the metabolic rate of organisms and previous studies have observed changes in enzymes involved in ATP production under anaerobic conditions (i.e., lactate dehydrogenase (LDH)) as well as enzymes involved in ATP production under aerobic conditions (i.e., citrate synthase (CS)) (Ransberry et al., 2015; Sifi and Soltani, 2019). Both of these metabolic enzymes, LDH and CS, have been used as biomarkers of the general physiological condition and used to evaluate the impacts of oxidative stress on organisms (Ransberry et al., 2015; Sifi and Soltani, 2019). Likewise, manganese superoxide dismutase (MnSOD) can provide another measure for estimating responses to hypoxic environmental conditions (Veldhoen et al., 2009). Prolonged hypoxia could induce the production of reactive oxygen species (ROS) leading to cellular damage, oxidative stress, and impaired energy production (Veldhoen et al., 2009). Accordingly, MnSOD acts as an antioxidant, recruited to inhibit the production of ROS (Meistertzheim et al., 2007).

Wastewater contamination has also been shown to induce the xenobiotic response in invertebrates and typically acts in a phased response pattern (Lüdeking and Köhler, 2002). The first phase (phase I) involves biotransformation enzymes like Cytochrome P450’s (e.g., CYP1A1) that modify contaminants by adding a functional group ensuring that the xenobiotics are more hydrophilic and are more easily biotransformed by phase II enzymes (Goksøyr, 1995; Morel et al., 1999). Phase II involves biotransformation enzymes like glutathione-S-transferases (GST) and glutamine synthetase (GS) that catalyze the binding of xenobiotics with endogenous substrates. The GST enzymes are highly conserved and not specific in binding electrophilic xenobiotics and organic contaminants, while GS specifically binds to nitrogen and ammonia and is highly regulated during xenobiotic exposure (Bao et al. 2013; Bonnafé et al. 2015). The final phase III biotransformation is crucial to the excretion of xenobiotics and prevents xenobiotics from entering the cytoplasm of a cell (Lüdeking and Köhler 2002; Bonnafé et al. 2015). Therefore, phase III transporters like the ATP-binding cassette (ABC) proteins and the multidrug resistance proteins (MDRP) are often used as biomarkers of exposure to xenobiotics (Lüdeking and Köhler 2002; Bonnafé et al. 2015). Changes in the expression patterns of these mRNA transcripts can be a result of a physiological response to an environmental stressor and subsequently, can provide a relative measure of contamination from the WWTP and act as an early warning system against prospective environmental changes (Etteieb et al., 2019; Kültz, 2005).

The results of climate change are causing people to aggregate in larger cities in northern regions, and as a result, there will be a greater impact of municipal wastewater effluent on Arctic marine systems in the future. In this study, we investigated the effects of primary treated municipal wastewater on the Arctic truncate soft-shell clam using a combination of endpoints including growth metrics, biogeochemical records (i.e., trace elements and stable isotopes), and mRNA expression profiles measured along a gradient in proximity to Iqaluit’s WWTP in Frobisher Bay, Nunavut, Canada. As the largest city in the Eastern Canadian Arctic, the impacts of wastewater effluent on marine organisms will likely be greatest near Iqaluit. The objective of this study was to determine the potential for the truncate soft-shell clam as a biomonitor for wastewater impacts in Arctic ecosystems by integrating long- and short-term physiological responses through sclerochronology and mRNA expression profiles. We hypothesized that continuous effluent from the WWTP would be a significant source of contamination to the Arctic ecosystem causing the truncate soft-shell clam to physiologically respond to the effluent, supporting its potential use as a biomonitor of wastewater in Northern Canada.

## Material and methods

### Sampling

A total of 292 *M. truncata* specimens were hand-collected from six sampling sites (Fig. in August 2019, using trowels during extreme low tide (<1 m) from Inner Frobisher Bay (63° 42’ 36’’ N, 68° 30’ 0’’ W). All collections were done with the support of the Amaruq Hunters and Trappers Association and under a license approved by Fisheries and Oceans Canada (Licence No: S-19/20-1040-NU). Each sampling site was chosen for its proximity to the WWTP (Fig. 1). Study sites included the WWTP, Tundra Ridge (3 km away from the WWTP), Apex (5 km away from the WWTP), Monument Island (5 km from the WWTP), Aupalajat (5 km away from the WWTP), and Kituriaqannigituq (15 km away from the WWTP). The WWTP collection site is located on the tidal flats and receives primary treated, year-round continuous municipal wastewater discharge. Aupalajat and Kituriaqannigituq were chosen as sites out of direct exposure from the wastewater outflow, protected by geographic barriers (i.e., Peterhead Inlet) and distance from the community (e.g., 15 km southwest), respectively.

**Figure 1.**
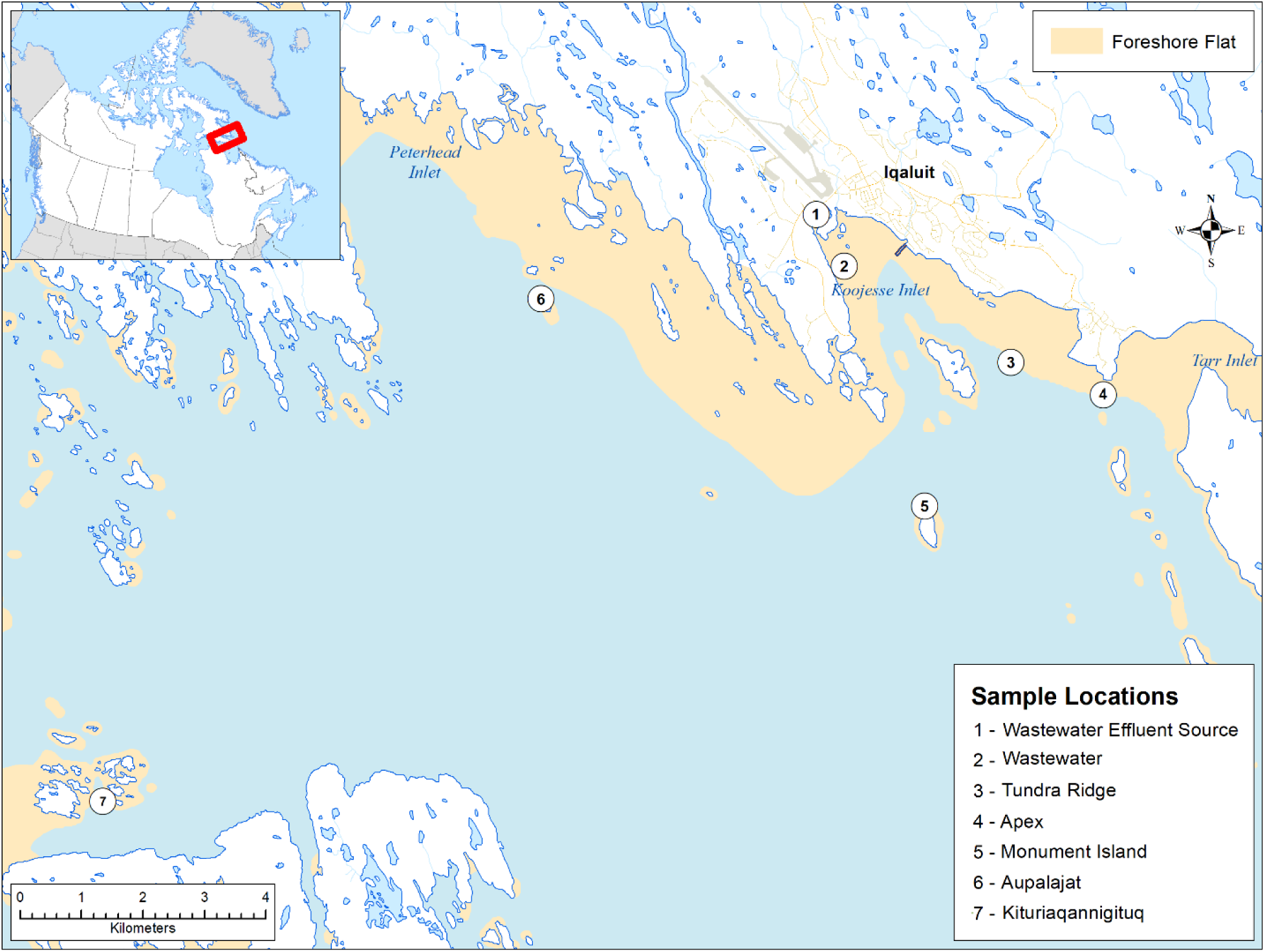
Location of Iqaluit’s wastewater effluent source and *Mya truncata* sampling locations in Inner Frobisher Bay, Nunavut, Canada. Inset shows area of study within Canada. This map was generated using ArcMap v10.7 with data from the National Atlas of Canada (National Atlas of Canada, 2020).

Shell length measurements were taken upon collection and were considered the maximum anterior – posterior dimension and measured with a pair of digital calipers (0.01mm; Mastercraft, Vonore, USA). Individuals chosen for sclerochronological sampling (n = 189) were placed in 4℃ temperature controlled 30L holding tanks for a maximum of 3 days with a recirculating artificial saltwater system at 100 gallons per hour before soft tissue was excised and shells were cleaned, air-dried, and weighed to the nearest gram (Instant Ocean Sea Salt; AquaClear Power Filter; Blacksburg, USA). These clams were held in tanks to allow for time sensitive mRNA samples to be processed. Seventy clams were also sampled for tissue specific mRNA patterns. Clams were opened by slicing their anterior and posterior adductor muscles and gill and mantle tissue were excised from each individual and immediately placed in 1 mL of RNA*later* (ThermoFisher, Waltham, USA). Tissues were stored and transported at −20℃ from Iqaluit, NU to the University of Manitoba, Winnipeg, MB where they were stored at −80℃.

### Growth pattern analysis

The age and internal growth bands in the shell were measured on the left chondrophore, a cavity that supports the internal hinge and yields the most distinct bands in *Mya* spp. An annual periodicity was assumed based on a previous study by MacDonald and Thomas (1980). Left shells were filled with epoxy resin (Varathane, Vernon Hills, USA) and sectioned from the umbo to the shell margin along the axis of maximal growth with a low-speed precision diamond saw continuously cooled in deionized water (Buehler, Uzwil, Switzerland). Once cut in half, the side of the valve containing the nucleus of the umbo was re-embedded in a thin layer of resin. The embedded halves were then ground and polished using a standard lapidary wheel (CrystalMaster, Green Meadows, USA) with a sequence of 350-, 600-, 1200-grit silicon carbide wet-table paper and finished with a polishing pad coated in 0.3 µm alumina suspension (MicroPolish, Buehler, Uzwil, Switzerland). The cross sections were etched and stained using a 100:100:1 mixture of 25% glutaraldehyde, 10% acetic acid, and amido black stain and left for 10 min before extracting the shells and rinsing three times in deionized water and air drying (Sigma Aldrich, St. Louis, USA).

To analyze the growth patterns, the chondrophores were imaged using a digital camera mounted on a dissecting microscope (Leica EZ4W, Wetzlar, Germany). The widths of the growth bands were measured digitally using the software Image J (Abràmoff et al., 2004). Band widths were measured independently by two people. Any disagreement between the two measurements was consistently due to pseudo growth bands grouped closely together. In the rare instance that both readers could not agree, the annual readings were discarded.

### Stable isotope analysis

Stable isotope analysis was performed on the two oldest specimens from each sampling location. Sample sizes (n = 2) were chosen to display intrashell variability that is lost with averaged values and replicate samples were taken within the individuals (n = 33-48). The periostracum was first removed using a rotary tool (Black & Decker, Towson, USA). Carbonate powder samples (50-100µg each) were then taken at intervals of 0.2-0.6 mm along the axis of maximal growth, using a dremel drill (Dremel^TM^, Wisconsin, USA) and later analyzed at the Manitoba Isotope Research Facility at the University of Manitoba. Stable oxygen (δ^18^O) and stable carbon (δ^13^C) isotopes were evaluated by a Costech^TM^ 4101 Element Analyzer (Costech Analytical Technologies Inc., Valencia, USA) coupled to a Thermo Finnigan^TM^ Delta V plus isotope-ratio mass spectrometer via an open split interface (ThermoFinnigan, San Jose, USA). Isotopes were normalized against calibrated international calcite standards, NBS-18 and −19 taken at the beginning, middle and end of each run. To check the quality of analysis performance, isotopes were calibrated against an internal calcite standard, and analysed together with unknown samples within this dataset (δ^18^O = ± 0.07‰; δ^13^C = ± 0.06‰). Shell results are reported in δ-notation for δ^18^O and δ^13^C and in per-mille (‰) versus VPDB (Vienna Pee Dee Belemnite Standard). Analytical precision (1σ) was ± 0.08‰ and ± 0.05‰ for δ^13^C and δ^18^O, respectively.

The δ^18^O ratio of seawater (δ^18^O_seawater_) was estimated using Ganssens equation and reported in ‰ versus VSMOW (Vienna Standard Mean Ocean Water; Witbaard et al., 1994). Salinity (psu) was sourced from multiple datasets across time (Supplementary Table S1) to reconstruct more reliable environmental conditions experienced by *M. truncata* due to scarce Arctic records (Bannister et al., 2013; Fisheries and Oceans Canada, 2019, 2018; Spares et al., 2012) and constants were based on cold water geographical regions (i.e., North Atlantic; Witbaard, 1997):

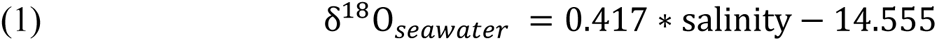

Seawater temperatures (℃) were then reconstructed from δ^18^O_shell_ (VPDB) and δ^18^O_seawater_ (VSMOW) values. Constants were defined based on all marine aragonitic molluscs and the subtraction of 0.2‰ adjusted the water and carbonate δ^18^O measurements to account for the fractionation between water and aragonite and the different scales δ^18^O was measured on (i.e., VPDB versus VSMOW). This modified equation was empirically derived for aragonitic bivalves and yields a temperature relationship precise to 1℃ accuracies in water greater than −10‰ (VSMOW) (Grossman and Ku, 1986):

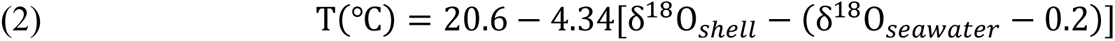

### Trace element analysis

Chondrophores from the right shell were mounted in 0.25-inch-thick epoxy mounts; ground and polished to obtain best analytical results. Chondrophores were cleaned in 10 min intervals to eliminate any external contamination using an ultrasonication cleaning bath (Bransonic Ultrasonic Cleaner; Brookfield, USA), an acetone bath (≥99.5% Acetone; Sigma-Aldrich, St. Louis, USA), a deionized water bath and finally another acetone bath, all conducted on a platform rocker (Vari-mix, ThermoFisher, Waltham, USA). These were left to dry overnight with filtered air passing over the samples.

Trace elemental concentrations – ^138^Ba, ^25^Mg, ^55^Mn, ^75^As, ^86^Sr, ^95^Mo, ^107^Ag, ^111^Cd, ^118^Sn, ^202^Hg, ^208^Pb, ^238^U, ^7^Li, ^23^Na, ^47^Ti, ^56^Fe, ^60^Ni, ^63^Cu, ^66^Zn – in chondrophores were analyzed using laser ablation inductively coupled plasma mass spectrometry (LA ICP-MS) on a New Wave UP-213 laser ablation system (Merchantek, Frement, USA) connected to an Element 2 high resolution ICP-MS (ThermoFinnigan, San Jose, USA). A total of 46 chondrophores (n = 7-8 chondrophores per location) were analyzed over 15 analytical sessions using both medium and low frequencies. Each ablation chamber held three resin-embedded chondrophores from different locations to avoid potential batch effects and machine drift for each sample series. The United States Geological Survey (USGS) carbonate reference material MACS-3 was ablated in duplicate and used as an external calibrator for marine carbonate-like minor elements (Pearce et al., 1997). A secondary external reference, NIST610, was ablated in duplicate to check quality and monitor machine drift (Pearce et al., 1997). MACS-3 and NIST610 were run at the beginning of each ablation chamber sample change, allowing for any correction during analysis. Specifications concerning LA-ICP-MS parameters and precision/accuracy can be found as a supplementary table (Supplementary Table S2). Data reduction (i.e., mapping, subtracting background, calculations of concentrations and limit of detection (LOD)) used *Igor Pro* graphing software (Wavemetrics Incorporated, Portland, USA) with an Iolite v3.7 package for LA ICP-MS following procedures described by Paton et al. (2011). Background signatures were subtracted, and calcium (^43^Ca) counts per second were used as an internal standard that normalized each element. Those elements with values that fell below the LOD or equal to below zero, were eliminated. By recording the start time, location and direction of the trace element scans, we were able to calibrate elemental results to calendar years within each growth line using ImageJ (Abràmoff et al., 2004).

### mRNA expression

Total RNA was isolated from the gill and mantle tissues using the RNeasy Plus Mini Prep Kit following manufacturer’s protocols (Qiagen, Hilden, Germany). Total RNA concentration and purity were assessed using a NanoDrop One spectrophotometer (ThermoFisher, Waltham, USA) and RNA integrity was further examined by gel electrophoresis. Total RNA samples were stored at −80℃ until further use. All samples were diluted to 1 µg of total RNA and were used to synthesize cDNA using a QuantiTect Reverse Transcription Kit (Qiagen, Hilden, Germany) in accordance with manufacturer protocols. cDNA was stored at - 20℃ until gene expression analysis.

A total of ten candidate genes that may represent a cellular response to wastewater effluent and two reference genes were selected for gene expression analysis (Table 1). Gene-specific primers for unique exon regions were designed from sequences from the assembled reference transcriptome (Sleight et al. 2018) for *M. truncata* on the mollusc database (https://ensembl.molluscdb.org/index.html). Primer Express v3.0.1 (ThermoFisher, Waltham, USA) was then used to produce single amplicons with a size greater than 100 bp, an annealing temperature of approximately 60℃, and a GC content of approximately 56%. Forward and reverse primers were synthesized by Integrated DNA Technologies (IDT; Coralville, IA, USA).

**Table 1.**
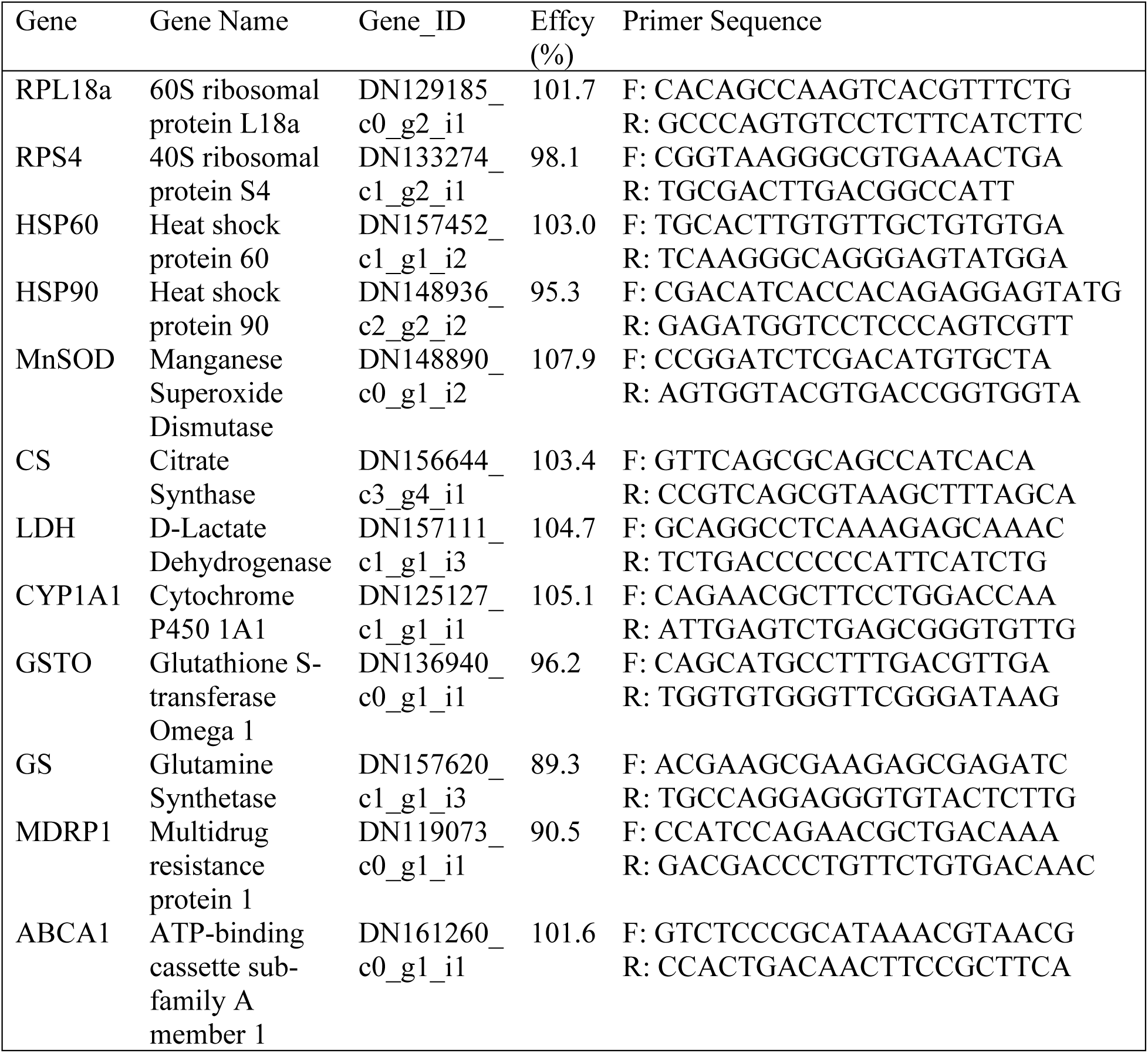
Primer sequences used to quantify mRNA transcript abundance of genes associated with stress, metabolism, and contaminant exposure in *Mya truncata* using qPCR. The Gene_ID’s are from the assembled reference transcriptome in Sleight et al. (2018) and the sequences were downloaded from the MolluscDB (https://ensembl.molluscdb.org/index.html).

Primer efficiencies (90-110% accepted) were assessed by using a 1:5 dilution standard curve for each primer set using pooled cDNA (Table 1). For a single reaction, primer mix was made using 0.5 µL of forward and reverse primers diluted to a concentration of 60 µM each using UltraPure water (ThermoFisher, Waltham, USA). This primer mix was further diluted with 0.9 µL of UltraPure water then combined with 6 µL of PowerUp™ SYBR™ Green (ThermoFisher, Waltham, USA) following the manufacturer’s guidelines, together making the PCR master mix. All qPCR reactions were completed using a QuantStudio 5 Real-Time PCR System (ThermoFisher, Waltham, USA) on 384 well-plates and were run under the same two-step cycling conditions: 2 min at 50℃, 2 min at 95℃, followed by 40 cycles of 15 s at 95℃, 15 s at 60℃ and 1 min at 72℃. Melt curve parameters consisted of denaturation for 15 s at 95℃ and 1 min decrease to 60℃ changing at a rate of 1.6℃ s^-1^, followed by a gradual increase to 95℃ for 15 s rising at a rate of 0.15℃ s^-1^. Twelve genes were run over three 384-well plates for each tissue and each plate included no template controls.

All expression data were normalized to the two reference genes, RPL18a and RPS4. These reference genes showed no significant differences across sampling locations and were considered stable across treatments when the cycle threshold (Ct) coefficient of variation was less than 25% (Hellemans et al., 2007). Relative changes in the expression of genes of interest were analyzed based on the 2^-ΔΔCt^ method (Livak and Schmittgen, 2001).

### Statistical analysis

#### Growth Pattern Analysis

The ontogenetic growth patterns displayed by the truncate soft-shell clams were detrended and standardized for each sample location. Upon comparing four common dendrochronology methods: smoothing spline, modified negative exponential (NE) curve, simple horizontal line or the modified Hugershoff curve; the detrending method with the lowest coefficient of variation within the timeseries was the negative exponential curve (Bunn, 2008). Thus, the primary ontogenetic growth signal was removed by applying an NE detrending curve:

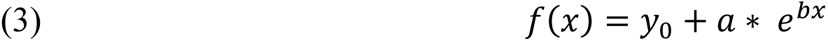

Where *f(x)* is shell length (mm), *y*_0_ is the shell length at *f*(∞), *x* represents the age (year), *a* is the intercept or length at first year of life, and *b* is the slope or rate of change in shell growth. *y*_0_, *a* and *b* are empirical constants which must be calculated for each species (Bunn, 2008). Any years that had less than three growth measurements were eliminated from this data set. Slopes (*b*) from the NE curves were then statistically compared between sampling locations using the R package emmeans (Searle et al., 1980) with a one-way analysis of variance (ANOVA) and pairwise comparisons were conducted using a Tukey’s Honest Significant Difference (HSD) post-hoc test. All growth standardization and analyses were conducted using the dendrochronology R package, dplR (Bunn, 2008).

#### Stable Isotopes

Isotope replicates were pooled for each shell and a Pillai’s Trace one-way multivariate analysis of variance (MANOVA) was conducted to determine the effects of sampling location on δ^18^O and δ^13^C stable isotope ratios. Follow-up univariate ANOVAs were used to assess if there was a significant difference in δ^18^O and δ^13^C between sampling locations. Individual mean comparisons across all six sampling sites and between δ^18^O and δ^13^C were analysed using Tukey’s HSD post-hoc test. The sea water temperature derived from δ^18^O (VSMOW; Equation was analysed with a one-way ANOVA and significance between locations was determined using Tukey’s HSD post-hoc test.

#### Trace elements

A multivariate principal components analysis (PCA) was conducted on the trace element data between sample locations and years to examine overall spatial and temporal trends. Differences between PC individual scores were statistically compared between location and year by a two-way ANOVA and multiple pairwise comparisons was computed by Tukey’s HSD post-hoc test. All multivariate analyses were performed using the R package FactorMineR (Lê et al., 2008). Furthermore, a cross-correlation function (CCF) was computed to assess the relationships (r) between growth rate and specific trace element signatures, Mn, and Sr. Mn and Sr are typically sensitive to biological influences like biomineralization rather than anthropogenic contamination sources. Thus, the CCF function was utilized to determine if their rate of uptake was dictated by ontogeny (Carré et al., 2006). This CCF function identifies how far the two time series are offset or ‘lagged’ (i.e., the number of years that naturally separate their correlation).

#### mRNA expression

Differences in relative mRNA abundance between sample locations were evaluated using a one-way ANOVA, followed by Tukey’s HSD post-hoc test for multiple comparisons for each tissue. A PCA was conducted to visualize the overall relationships between mRNA expression and sampling locations for each tissue type using FactorMineR (Lê et al. 2008). One-way ANOVAs were used to detect significant differences in individual PCA scores between sample locations and multiple pairwise comparisons were assessed using Tukey’s HSD post-hoc tests. All statistical analyses were assessed with the level of significance (α) of 0.05 and analyses were performed using the statistical computing software, R v3.6.3 (R Core Team, 2020).

## Results

### Analysis of Growth Variations

*M. truncata* specimens ranged from 4-32 years of age and between 26-90 mm in length (maximum anterior to posterior length). Sample size began to decline at eight years of age on the NE detrending curve and only eight specimens reached between 24 to 32 years of age (Fig. 2). *M. truncata* displays an ontogenetic growth pattern, where the growth rate declines with age following the shape of an exponential decay curve (Table 2; Román-González et al., 2017). This rate of change (*b*) in shell growth demonstrated significant differences between locations (one-way ANOVA: *F*_5,105_ = 82.6, *P* < 0.001; Fig. 2). Comparisons between groups revealed that the shell growth rate at the WWTP was significantly slower than at Apex, Monument Island, and Kituriaqannigituq (TukeyHSD: *b* = −0.01, *P <* 0.05; Fig. 2). Tundra Ridge also displayed significantly slower growth rates than Apex (TukeyHSD: *b* = −0.02, *P* < 0.05; Fig. 2). Data on the standardized growth index are provided within the supplementary figures (Supplementary Fig. S1).

**Figure 2.**
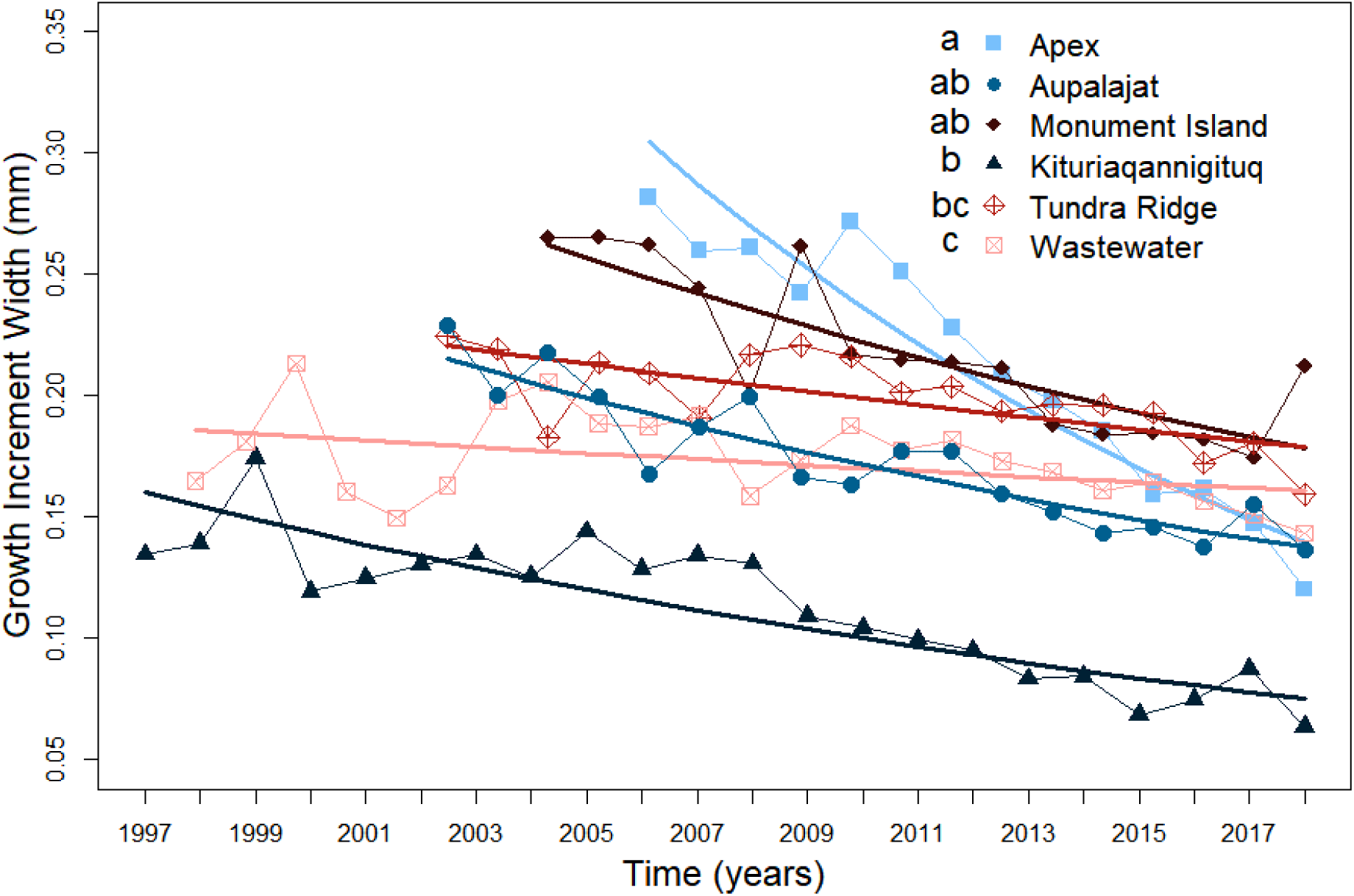
Raw ontogenetic shell growth (mm) over time (years) with detrending negative exponential curves for *Mya truncata* organisms in each of the six sampling locations in Inner Frobisher Bay, Nunavut, Canada. NE detrending curve equations can be found in Table 2 and follows an exponential decay pattern with y = 0 as an asymptote. Lowercase letters denote statistical significance between the slopes of sampling sites (α < 0.05) as determined by a one-way ANOVA and Tukey’s HSD post-hoc test.

**Table 2.** Resolved negative exponential detrending curve equations for *Mya truncata* shell growth (mm) at six sampling locations in Inner Frobisher Bay, Nunavut, Canada (Equation 2.3). CV represents the coefficient of variation.

| Site | Intercept (*a*) | Slope (*b*) | R<sup>2</sup> value | F-statistic | CV (%) |
| --- | --- | --- | --- | --- | --- |
| Apex | 118.88 | -0.06 | 0.90 | 117.1 | 8.7 |
| Aupalajat | 32.6 | -0.04 | 0.83 | 80.46 | 10 |
| Monument Island | 66.2 | -0.03 | 0.70 | 33.37 | 15.6 |
| Kituriaqannigituq | 59.4 | -0.03 | 0.69 | 49.97 | 3.2 |
| Tundra Ridge | 31.6 | -0.02 | 0.49 | 15.47 | 6.8 |
| Wastewater | 16.6 | -0.01 | 0.18 | 4.667 | 13.2 |

### Stable Isotopes Signatures

The average shell stable oxygen isotope profiles (δ^18^O) from each site ranged from 1.2-1.8‰ and the average shell stable carbon isotope profiles (δ^13^C) from each site ranged from 1.3-2.1‰ (Table 3). Mean isotope ratios were significantly different among the six sample locations (Pillai’s trace MANOVA: *F*_10,268_ = 12.5, *P* < 0.001; Fig. 3) and follow up univariate analysis displayed significant differences in δ^18^O (univariate ANOVA: *F*_5,163_ = 70.7, *P* < 0.001) and δ^13^C (univariate ANOVA: *F*_5,163_ = 70.2, *P* <0.001) between sampling locations. The clams near the WWTP had a distinct covariation with δ^13^C and δ^18^O ratios, demonstrating significantly lower isotopic ratios than any other site (TukeyHSD: *P* < 0.001; Fig. 3). A hierarchy formed with δ^13^C ratios displaying: Monument Island and Tundra Ridge >Apex and Aupalajat > WWTP, and Kituriaqannigituq showing no significant differences from any of the sites (Fig. 3).

**Figure 3.**
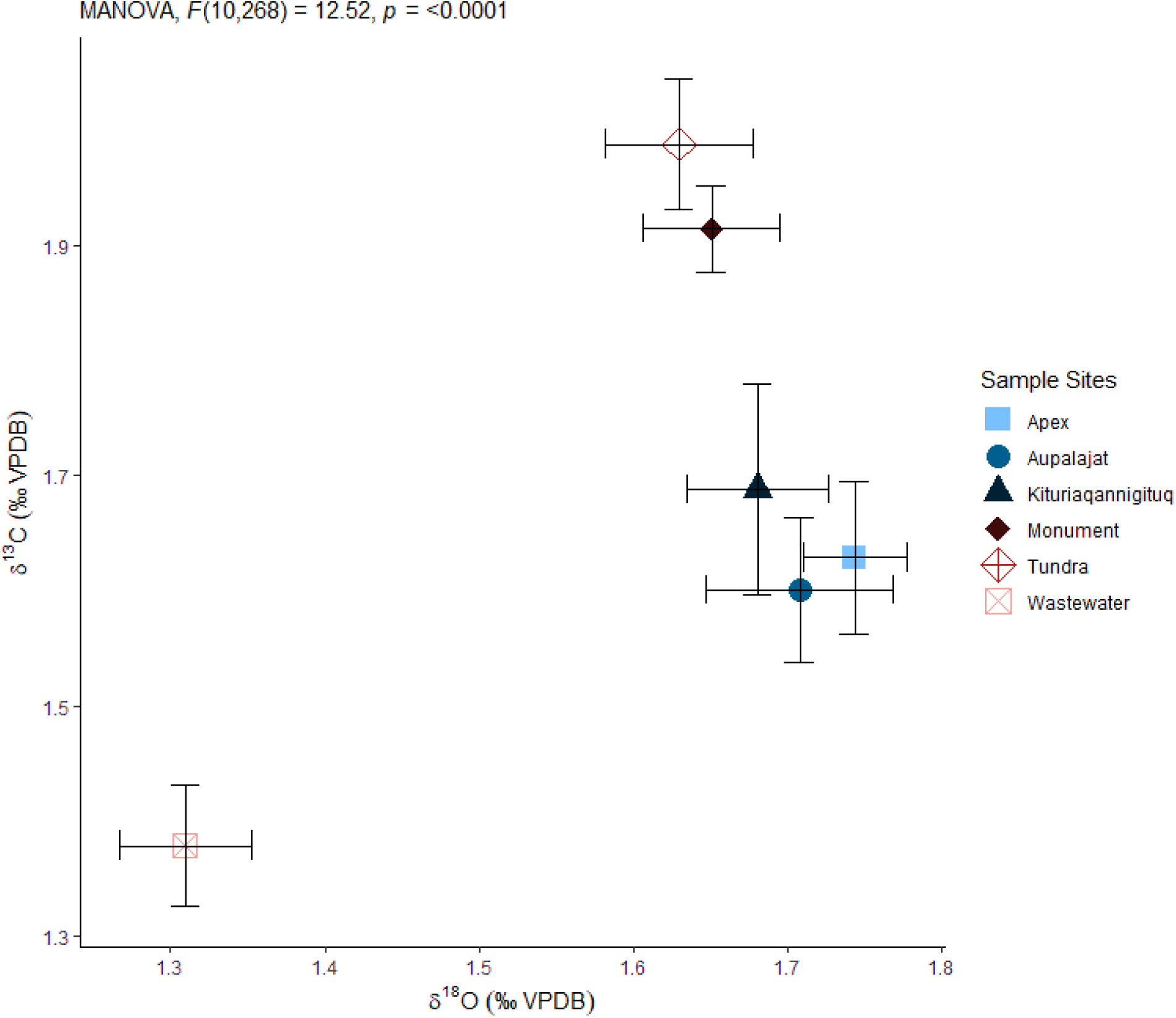
Average *Mya truncata* shell δ^18^O versus δ^13^C isotope ratios (‰ VPDB ± s.e.m.) for each respective sampling location in Inner Frobisher Bay, Nunavut, Canada. One-way MANOVA and Tukey’s HSD post-hoc tests were conducted to determine effect of sampling location on δ^18^O and δ^13^C stable isotopes (α < 0.05).

**Table 3.** Summary of stable isotope parameters from *Mya truncata* shells for each sample location in Inner Frobisher Bay, Nunavut, Canada. For each location, n represents the number of samples analyzed and s.e.m. is the standard error to the mean.

| Location | δ<sup>13</sup>C (n) | δ<sup>18</sup>O (n) | δ<sup>13</sup>C mean ± s.e.m | δ<sup>18</sup>O mean ± s.e.m. |
| --- | --- | --- | --- | --- |
| Tundra Ridge | 29 | 30 | 1.98 ± 0.06 | 1.65 ± 0.04 |
| Monument island | 26 | 22 | 1.92 ± 0.04 | 1.65 ± 0.04 |
| Aupalajat | 35 | 33 | 1.71 ± 0.05 | 1.81 ± 0.05 |
| Kituriaqannigituq | 24 | 23 | 1.70 ± 0.09 | 1.66 ± 0.04 |
| Apex | 25 | 25 | 1.63 ± 0.07 | 1.74 ± 0.03 |
| Wastewater plant | 33 | 33 | 1.38 ± 0.05 | 1.31 ± 0.04 |

Using shell δ^18^O (VPDB) and estimated seawater δ^18^O (VSMOW) (Equation 2), we found a statistically significant difference (one-way ANOVA *F*_5,165_ =13.66, *P* < 0.001; Supplementary Fig. S2) in reconstructed average temperatures (℃) at each location: WWTP (0.62 ± 0.20), Tundra Ridge (−0.68 ± 0.20), Apex (−1.17 ±0.15), Monument Island (−0.39 ± 0.27), Aupalajat (−1.63 ± 0.26) and Kituriaqannigituq (−0.90 ± 0.20). Pairwise comparisons determined that the WWTP (TukeyHSD: *P* < 0.01) had significantly higher temperatures than all other sites and Aupalajat was significantly lower than all other locations (TukeyHSD: *P* < 0.05).

### Trace Element Profiles

Generally, most elements exhibited significant patterns between sampling locations and time, but Mo, Ag, Cd, Sn, Li, and Zn did not differ temporally nor spatially and therefore, only those exhibiting significant patterns will be discussed herein. The first two dimensions of the PCA for the entire set of trace elements explained 37% of the total variance (Fig. 4). The variables driving separation of sites on PC1 (19.5% variance explained) were Cu, Pb, Mg, and Fe ranked in decreasing order of effect. Among individual profiles, these four elements consistently displayed higher overall concentrations at the WWTP and Tundra Ridge and the clams from these sites were on the positive end of PC1. Clams from Monument Island were on the negative side of PC1 with Aupalajat, Apex, and Kituriaqannigituq clustering in the middle. PC1 explained significant differences between locations but not between years (two-way ANOVA: *F*_5, 91_= 30.7, *P* < 0.001; Fig. 4). Arsenic (As), Sr, Mn, Ba, and Na variables (ranked highest to lowest in their effect) were responsible for the spatial (two-way ANOVA: *F*_5,91_ = 7.7, *P* < 0.001; Fig. 4) and temporal (two-way ANOVA: *F*_19,83_ = 18.3, *P* < 0.001; Fig. 4) separation on PC2 (17.5% variance explained). Temporally, seven groups were formed, showing a ranked chronology with year 2000 on the positive side of PC2, to year 2019 being on the negative side of PC2 (TukeyHSD: *P* < 0.05; Fig. 4). Along PC2, sample locations were split into two groups, with Monument Island, Apex, Aupalajat, and Kituriaqannigituq on the negative end of PC2 while Tundra Ridge and the WWTP were on the positive end of PC2 (TukeyHSD: *P* < 0.05; Fig. 4).

**Figure 4.**
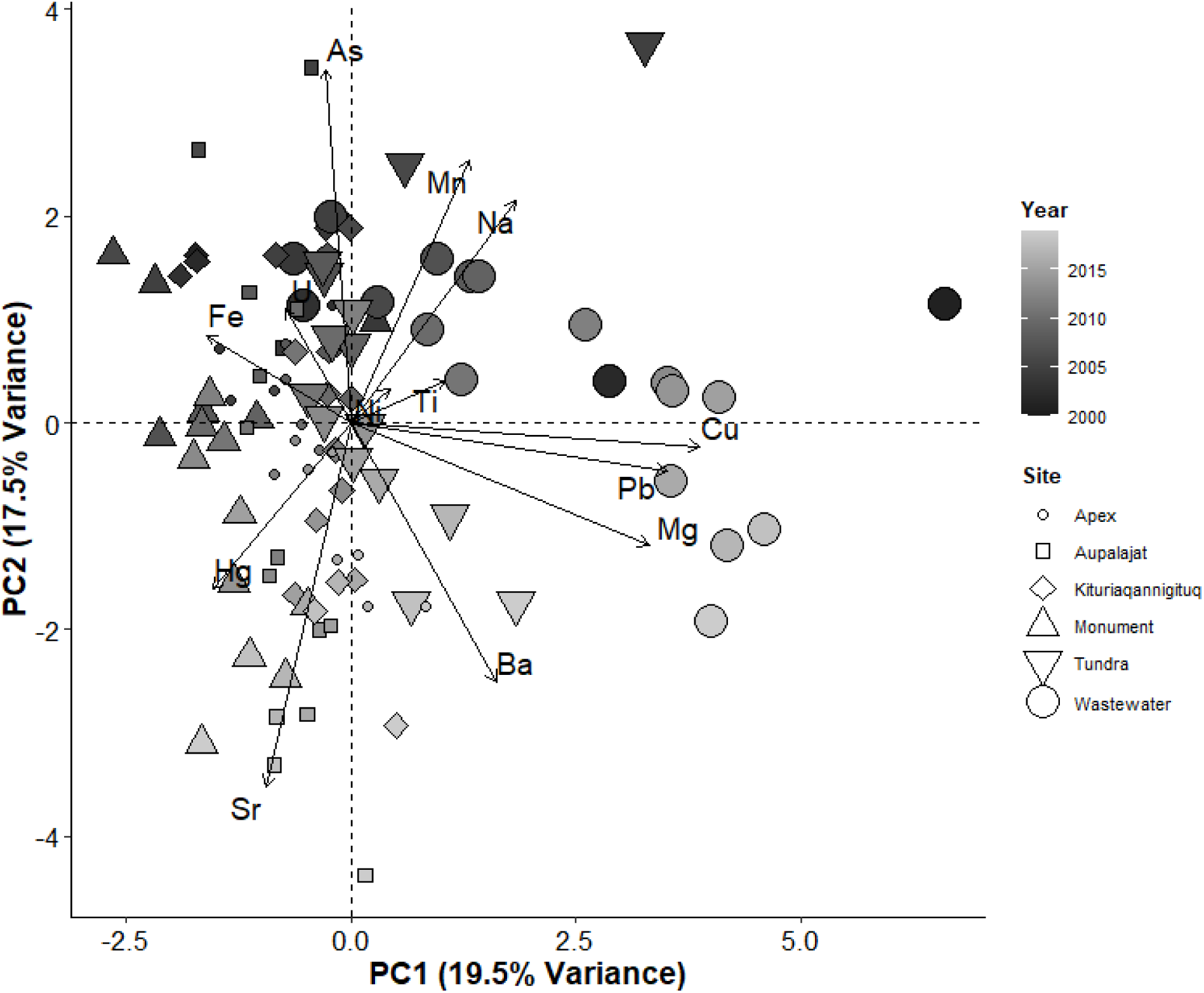
Principal components analysis biplot of trace elements found in the shells of *Mya truncata* organisms characterized by year and location. Years range from 2000 – 2019 and locations include the six sampling locations within Inner Frobisher Bay, Nunavut, Canada. X-axis represents PC1 (19.5% variance) while the Y-axis represents PC2 (17.5% variance).

The relationship between growth rate, Sr, and Mn were compared and these two elements followed opposite patterns over time as Sr had a marked increase in concentration and Mn had a decline in concentration over time within each sampling location. Cross-correlation analysis found a significant negative relationship between Sr and growth rate at all sites (CCF: r = −0.42 to −0.81, *P* < 0.05; Table 4) except the WWTP (CCF: r = 0.37; Table 4). Similarly, Mn displayed a positive relationship with growth rate at all sites (CCF: r = 0.57 to 0.78, *P* < 0.05; Table 4) except the WWTP which displayed a negative relationship (CCF: r = −0.16; Table 4).

**Table 4.** Pearson cross correlation values (r) and significance between strontium (Sr), manganese (Mn), and growth rate (GR) for *Mya truncata* shells in each sampling location in Inner Frobisher Bay, Nunavut, Canada. Lag represents the number of years that naturally separate the two time series (α < 0.05) and n.s. indicates no significance found.

| Site | Relationship | r | Lag (yrs) | p-value |
| --- | --- | --- | --- | --- |
| Aupalajat | Sr:GR | -0.69 | 1 | <0.05 |
|  | Mn:GR | 0.78 | 0 | <0.01 |
| Kituriaqannigituq | Sr:GR | -0.42 | 0 | n.s. |
|  | Mn:GR | 0.68 | 0 | <0.05 |
| Wastewater | Sr:GR | 0.37 | -4 | n.s. |
|  | Mn:GR | -0.16 | -9 | n.s. |
| Tundra Ridge | Sr:GR | -0.81 | 0 | <0.01 |
|  | Mn:GR | 0.57 | -2 | <0.05 |
| Monument Island | Sr:GR | -0.65 | -1 | <0.05 |
|  | Mn:GR | 0.63 | -1 | <0.05 |
| Apex | Sr:GR | -0.69 | -1 | <0.01 |
|  | Mn:GR | 0.7 | -2 | <0.05 |

### mRNA Expression Profiles

#### Gill mRNA Abundance

We identified whether the abundance of select genes were significantly different among the six sampling locations. The first two PC dimensions explained 45.4% of total variance in the overall gill expression among sampling locations (Fig. 5). PC1 (29.1% variance explained) was driven by the cellular stress response genes GS, HSP90, GST, and LDH and the phase III xenobiotic response ABCA1 (one-way ANOVA: *F*_5,51_ = 4.6, *P* < 0.01; Fig. 5). The abundance of all five mRNA transcripts were low at the WWTP and Kituriaqannigituq and elevated at Aupalajat with varying degrees of expression (TukeyHSD: *P* < 0.05; Supplementary Fig. S3). Due to these similar expression patterns, PC1 had Kituriaqannigituq, the WWTP, Apex, and Tundra Ridge on the negative side of the PC1 axis and Aupalajat on the positive side of the PC1 axis (TukeyHSD: *P* < 0.05; Fig. 5). Monument Island individuals were mixed between the two groups. PC2 (16.3% variance explained) was primarily driven by genes MDRP1 and MnSOD (one-way ANOVA: *F*_5,51_ = 5.0, *P* < 0.001; Fig. 5). MnSOD displayed a similar expression pattern with Monument Island and the WWTP having low abundance but Kituriaqannigituq having the highest expression (TukeyHSD: *P* < 0.01; Supplementary Fig. S3). MDRP1 exhibited significantly different abundance among sampling locations (one-way ANOVA: *F*_5,51_ = 5.3, *P* < 0.01; Supplementary Fig. S3) with a 3-fold increase in expression at the WWTP compared to all other locations (TukeyHSD: *P* < 0.001; Supplementary Fig. S3). Three groups formed in pairwise analysis of PC2 significantly separating the WWTP on the negative side of the PC2 axis and Kituriaqannigituq on the positive side of the PC2 axis (TukeyHSD: *P* < 0.001; Fig. 5).

**Figure 5.**
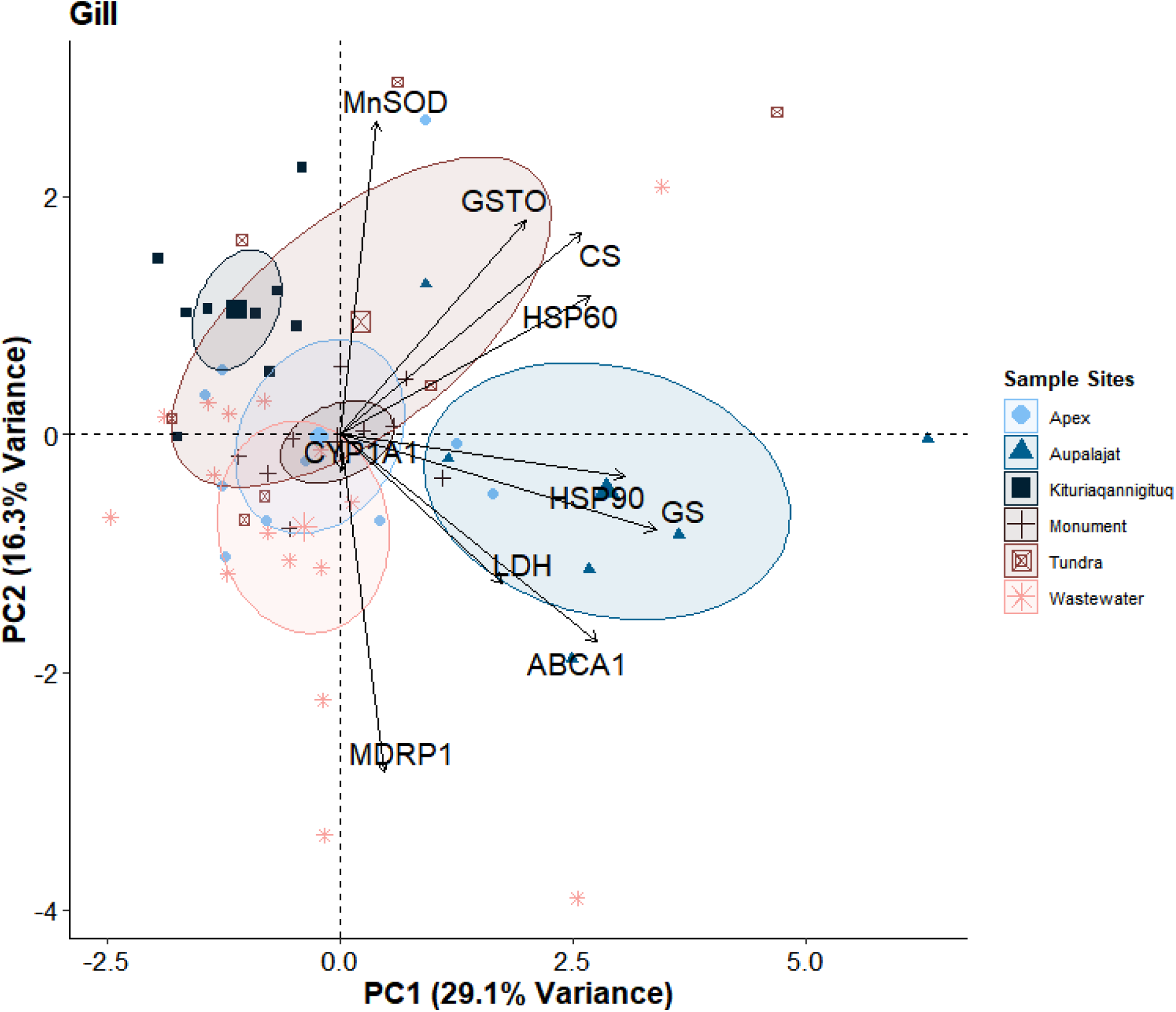
Principal components biplot analysis on the gill gene expression for *Mya truncata* organisms from the six sampling locations in Inner Frobisher Bay, Nunavut, Canada. The X-axis represents PC1 (29.1% variance), while the Y-axis represents PC2 (16.3% variance). Larger symbols indicate the mean for that sample site.

#### Mantle mRNA Abundance

Mantle tissues examined had a lower abundance of mRNA and displayed less significance in expression among sampling locations than the gill. The first two PC axes explained 41.9% of the total variance in genes expressed in the mantle among sampling locations (Fig. 6). The significant separation between sampling location on PC1 (24.4% variance explained) was primarily driven by the cellular stress response gene LDH and phase II xenobiotic response gene GS (one-way ANOVA: *F*_5,50_ = 6.8, *P* < 0.001; Fig. 6). LDH expression was highest at Apex and lowest at Kituriaqannigituq (TukeyHSD: *P* < 0.05; Supplementary Fig. S4). Similarly, the highest abundance of GS was displayed at the WWTP, Apex, and Aupalajat and had the lowest expression at Kituriaqannigituq (TukeyHSD: *P* < 0.05; Supplementary Fig. S4). These significant differences led to Kituriaqannigituq separating on the negative end of PC1 and Apex, the WWTP, and Aupalajat on the positive side of PC1 (TukeyHSD: *P* < 0.05; Fig. 6). The PC2 axis (17.5% variance explained) was primarily driven by the genes CYP1A1 and MDRP1 and while these genes showed significant patterns between sample locations, there was no significant separation of sites along PC2. The phase III biotransformation gene MDRP1 was highly expressed at the WWTP and exhibited a gradual decline in expression with increasing distance from the wastewater effluent source having its lowest abundance at Monument Island, Kituriaqannigituq, and Aupalajat (one-way ANOVA: *F*_5,41_ = 3.4, *P* < 0.05; Supplementary Fig. S4).

**Figure 6.**
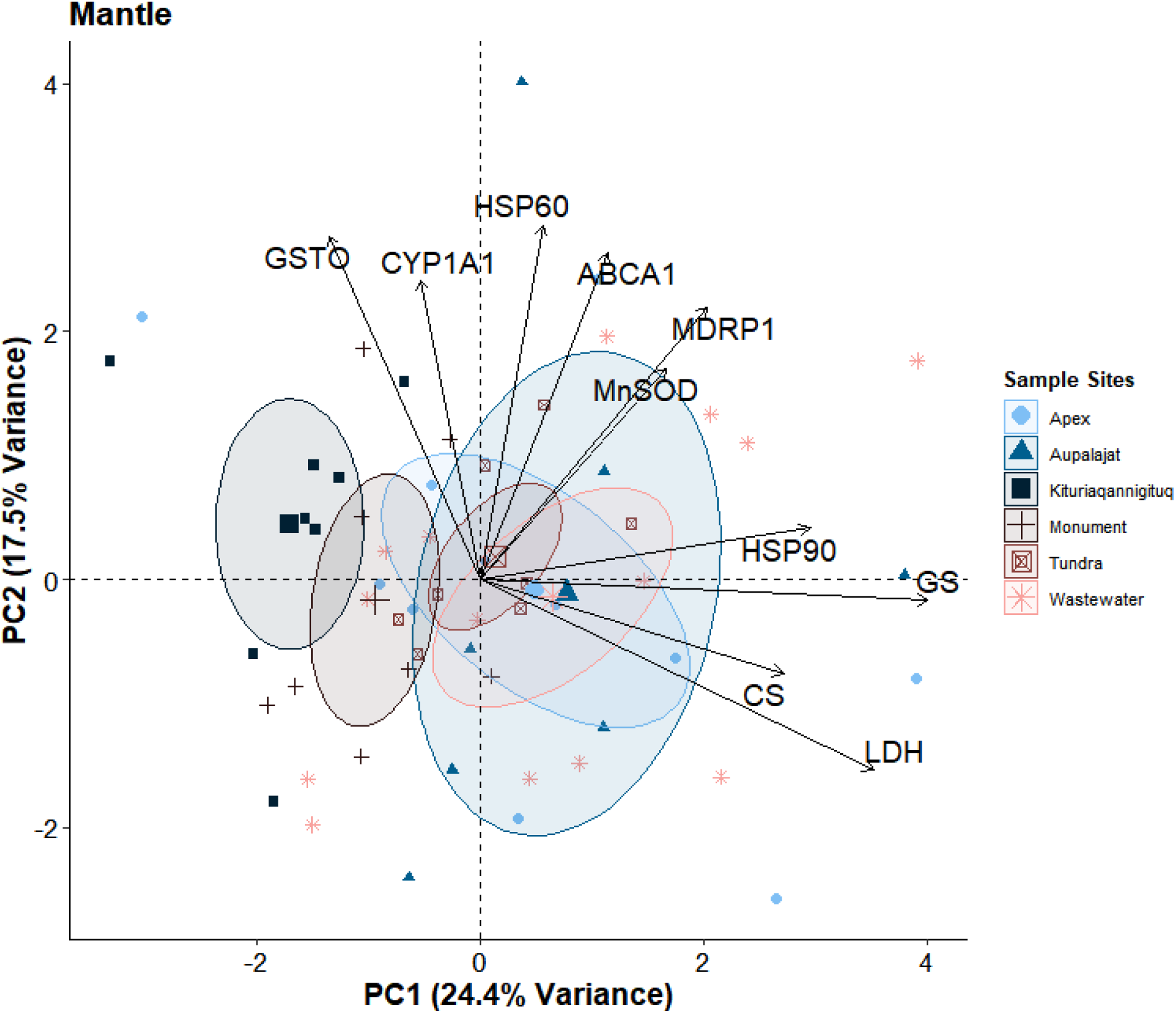
Principal components biplot analysis on the mantle gene expression for *Mya truncata* organisms from the six sampling locations in Inner Frobisher Bay, Nunavut, Canada. The X-axis represents PC1 (24.4% variance), while the Y-axis represents PC2 (17.5% variance). Larger symbols indicate the mean for that sample site.

## Discussion

This study explicitly examined the effects of municipal wastewater on the Arctic truncate soft-shell clam using a combination of endpoints including growth metrics, biogeochemical records (i.e., trace elements and stable isotopes), and mRNA abundance profiles to interpret the impacts of wastewater contamination in Inner Frobisher Bay. The municipal wastewater treatment plant in Iqaluit discharges approximately 1.2 x 10^6^ m^3^ of municipal wastewater into Frobisher Bay each year (Neudorf et al., 2017). This volume is set to increase as Iqaluit has seen a dramatic population increase, rising 15.6% between 2011 and 2016 (Statistics Canada, 2017). The raw effluent consistently exceeds the federally regulated maximum for biological oxygen demand (BOD), total suspended solids (TSS), total and fecal coliform concentrations, and numerous studies have documented the presence of ammonia (City of Iqaluit, 2019), phosphate, phosphorus, and metals (e.g., Al, Ba, Cd, Cu, Fe, Mn, Hg, Ag, Sr, and Zn; Daley et al., 2015; Krumhansl et al., 2015; Nunami Stantec, 2015). The impact of this discharge on organisms in the Arctic environment is largely unknown. We found evidence of reduced growth rate, elemental accumulation in the shell, and covariant depleted stable isotopes in clams sampled near the WWTP suggesting that the wastewater effluent is a source of contamination to the aquatic environment. Furthermore, detailed sclerochronological record of isotopes and elements substantiated the truncate soft-shell clam’s potential to be an Arctic biomonitor species. Finally, the decrease in abundance of mRNA transcripts of molecular chaperones, antioxidants, metabolic biomarkers, and phase I and II biotransformation enzymes concurrently happened alongside an increase in the expression of phase III xenobiotic defense genes and the nitrogen specific binding enzyme, which suggested that these bivalves were physiologically responding to the chronic contamination source.

### Growth Metrics and Stable Isotopes

Growth rate is a common metric used to assess physiological responses to contamination sources in bivalves (Nobles and Zhang, 2015; Sabatini et al., 2011). Previous studies have found growth impairment in bivalves upon chronic exposure to wastewater effluent sources in a wide array of environments, using diverse wastewater treatment methods (i.e., primary to tertiary; Gangloff et al., 2009; Goudreau et al., 1993; Nobles and Zhang, 2015). The present study agrees with these previous observations as the relative growth rate of the truncate soft-shell clam was significantly lower at the two sites nearest the WWTP (i.e., WWTP and Tundra Ridge) compared to more distant locations. Krumhansl et al. (2015) also found altered community structure in benthic fauna 580 m away from the WWTP in Iqaluit. Benthic sediments were anoxic due to nutrient enrichment and had visible reductions in species diversity, density, richness, and evenness. Our study builds on these findings, showing the alteration of softshell clam physiology and growth nearly 3 km away at Tundra Ridge, highlighting concerns for the range of impact from the WWTP. Numerous factors have been shown to impact the growth rate of bivalves, but the isotopic reconstruction of heightened temperature, lower salinity, and elevated organic matter confined to the WWTP site in our study supports the notion that these environmental variations are likely anthropogenically-driven (Brown, 1978; Schöne and Surge, 2012).

Elevated water temperatures are common in the receiving waters of wastewater effluent (Francy et al., 1996; Tuncay, 2016). Iqaluit’s effluent has previously been recorded to be as warm as 15℃ discharging into the ∼1.9℃ receiving waters (in 2013 and 2014 Neudorf et al., 2017). This supports the higher average reconstructed sea temperatures (via δ^18^O VSMOW) in this study and could influence the reduced growth in these organisms. While these organisms could achieve maximal growth within an ideal temperature range, minor increases beyond this range could depress these growth rates (Nobles and Zhang, 2015; Talmage and Gobler, 2011). The reconstructed sea temperatures only reflect ambient temperatures during active growth and therefore do not represent temperature for the entire year.

Many studies have recorded the utility of covarying trends of δ^13^C and δ^18^O recorded in bivalve shells and the depleted ratios found in the clams nearest the WWTP detected in the present study could be indicative of brackish water and increased organic matter (Jones and Quitmyer, 1996). The distinctive pattern of elevated organic matter near the wastewater outfall is further supported by the incorporation of Ba in the shell. Ba is often used to discern ocean productivity and nutrient dynamics, thus, its higher concentration paired with the reproducible seasonality at the WWTP, Apex, and Tundra Ridge provides a potential record of the WWTP discharging nutrient enriched effluent to the area (Putten et al., 2000; Stecher et al., 1996).

In the present study, the incorporation of Sr and Mn into the shell are under the strong biological influence of biomineralization rates rather than anthropogenic sources. Therefore, the rate of Sr and Mn elemental incorporation is dictated by ontogeny and metabolism. The rate of uptake is dependent on the size of ions as well as the organisms affinity for calcium uptake over other elements (Carré et al., 2006; Ferrier-Pagès et al., 2002). In this study, Sr showed strong negative correlations while Mn displayed strong positive correlations with growth rate at all sites except the WWTP. This irregular pattern from the WWTP could be explained by the overall slower growth rate or site-specific differences in sea temperature, nutrient availability, salinity, or toxicity.

### Trace Elements

The temporal progression of elements in the shells of individuals nearest the wastewater effluent source display the benefit of primary treatment by containing higher concentrations of Mn, Na, and Ti from 2000-2010 to incorporating higher concentrations of Cu, Pb, and Mg in later years, 2010-2018. These temporal trends can therefore potentially be attributed to two things. The first is the implementation of primary treatment in 2006, which included the installation of a Salsnes filter that eliminates elements already low in concentrations from being released into the receiving waters and has been shown to remove approximately 50% of TSS in wastewater effluent (Nunami Stantec, 2015). The increased concentration of Cu, Pb, and Mg between 2010 and 2018 may be a result of a rising population (15.6% increase; Statistics Canada, 2017) eliciting more effluent volume and thereby causing elements most common to wastewater discharge to be at higher concentrations. In general, metals like Pb and Cu do not show any predictable variance with respect to environmental and biological controls and hence their variability can be linked to local contamination sources (Price and Pearce, 1997).

### mRNA Expression Patterns

Given the evidence of metal accumulation and isotopic change in clams near the WWTP, we expected to see physiological responses to these contaminants and modifications in mRNA abundance related to the xenobiotic response (i.e., phase I, II and II biotransformation biomarkers) and cellular stress response (i.e., molecular chaperones, antioxidants, metabolic regulators). When exposed to contamination, xenobiotics have been shown to greatly enhance the abundance of CYP1A1 (Viarengo et al., 2007); an enzyme regularly recruited to ensure xenobiotics are more easily removed from the cell (Goksøyr, 1995). This study displayed opposite results as CYP1A1 had a lower expression near the WWTP. However, a negative feedback loop occurs in that as CYP1A1 activity increases, so does the production of its by-products, ROS and hydrogen peroxide (Morel et al., 1999). The resulting hydrogen peroxide may act to repress CYP1A1 at the transcriptional level to ultimately prevent ROS production and oxidative stress (Barouki and Morel, 2001; Morel and Barouki, 1998). This mechanism is a possible cause for the decreased CYP1A1 activity at the four sites nearest the wastewater effluent source relative to the two sites shielded from direct wastewater exposure (i.e., Aupalajat and Kituriaqannigituq).

The phase II biotransformation enzyme GST operates to bind xenobiotics by conjugating glutathione, acting as a first step towards their elimination from the cell (Bonnafé et al., 2015). The reduced abundance of GST mRNA transcripts near the WWTP is supported by numerous other studies on bivalves demonstrating a decrease in GST expression when exposed to chronic wastewater (Ballesteros et al., 2009; Bonnafé et al., 2015; Lüdeking and Köhler, 2002). One hypothesis suggests that this low expression is a result of excess metabolites produced by CYP1A1 competing with GST endogenous substrates (Bonnafé et al., 2015).

Another phase II enzyme in the xenobiotic response is GS, which specifically binds to nitrogen and ammonia and plays an essential role in the metabolism of nitrogen (Bao et al., 2013). GS binds to ammonia to synthesize glutamine, an amino acid heavily involved in the immune system of bivalves (Hanson and Dietz, 1976). Its increased expression has been linked to increased protein catabolism, amino acid turnover, nitrogen detoxification, and nitrogen pollution (Tanguy et al., 2005; Thomsen et al., 2016). We found greater expression of GS activity at the WWTP, Apex, and Aupalajat, which could be related to both nitrogen pollution and increased production of nitrogenous waste products. P-glycoproteins (P-gp) like MDRP1 and ABCA1 contribute to the removal of xenobiotics out of the cell through ATP-dependent pumps to prevent intracellular toxicity (Smital et al., 2000; Veldhoen et al., 2009). In both tissue types, MDRP1 was significantly upregulated at the WWTP, which may be due to the increased need to eliminate xenobiotics from the cell (Lüdeking and Köhler, 2002).

Heat shock proteins are used to measure a vast array of physiological stress in bivalves and more specifically, HSP90 is recruited to maintain homeostasis and protect the cell against xenobiotics (Fabbri et al., 2008). In this study, the low abundances of HSP90 in the gills of clams nearest the WWTP suggests that expression is more related to oxidative stress than other stimuli as chronic exposure to contamination could inhibit its function (Fabbri et al., 2008; Foster and Fulweiler, 2019). The conditions of oxidative stress at the WWTP is further supported by the expression pattern of MnSOD, a chief ROS scavenging enzyme (Holley et al., 2011). Excess contaminants and metal exposure have been shown to interfere with ROS defense mechanisms like MnSOD, especially in cold water environments (Cossu et al., 2000; Ransberry et al., 2015). Thus, the low abundance of MnSOD displayed at the WWTP is potentially a response to contamination. Interestingly, the Kituriaqannigituq reference site also had low expression of HSP90 and MnSOD, which suggests potential oxidative stress and hypoxic conditions in that region. Lactate dehydrogenase is another gene that could support the presence of hypoxic conditions and the potential impact on metabolic rates. The reduced LDH expression displayed in the present study could reflect a possible decrease in overall biosynthetic activities under chronic exposure conditions and ultimately result in the decreased capacity for anaerobic ATP production under hypoxic conditions (Sifi and Soltani, 2019; Sonawane, 2017). Further investigation on the environmental conditions and mitochondrial capacity of the truncate soft-shell clam is required, but an inhibition of LDH could indicate a reduction in metabolic capacity, potentially associating this physiological response to the reduced growth seen in soft-shell clams nearest the WWTP site.

## Conclusion

We present an empirical assessment of *Mya truncata* as a biomonitor of municipal wastewater in remote Arctic communities. As northern communities continue to grow, the ecological implications of chronic wastewater introductions to coastal marine systems will also increase. Over time, wastewater effluent could disrupt the ecological processes and physiological function of Arctic marine organisms. We show that the truncate soft-shell clam has already displayed modified physiological function in response to wastewater exposure near Iqaluit, the largest city in Eastern Arctic Canada. Having a relevant organism to assess the impact of contaminants on Arctic marine systems will be extremely useful for monitoring the effects of wastewater effluent in ecosystems where conventional monitoring programs are challenging.

## Acknowledgements

This project was supported by funds from Fisheries and Oceans Canada (DFO) Coastal Environmental Baseline Program, as well as an NSERC Discovery Grant (#05479) awarded to KMJ, and a University of Manitoba University Indigenous Research Program grant (UM project #50683) awarded to KMJ and DD. We thank the Amaruq Hunters and Trappers Association for their guidance and collection license. We also thank Christopher Lewis, Davidee Qaumariaq, and Alex Flaherty for their field assistance, coordination, and sample collection. Moreover, we thank Jordan Kroeker and Isabel Hilgendag for their field and sample collection assistance. We gratefully acknowledge Mark Ouellette for constructing the site map and Daniel Gedig for acting as a secondary ager for this study. Furthermore, we thank Victoria Sleight for providing annotated reference transcriptomes. We appreciate Panseok Yang for help with operating the LA ICP-MS and Misuk Yun for operating and conducting stable isotope analyses. We thank Dirk Weihrauch and Mark Hanson for input on analyses and interpretations, and Jason Treberg for the use of his microscope.

