## Supplementary Materials for "The truncate soft-shell clam, *Mya truncata*, as a biomonitor of municipal wastewater exposure and historical anthropogenic impacts in the Canadian Arctic"

Christina M Schaefer

- University of Manitoba, Winnipeg, Manitoba, Canada
- Fisheries and Oceans Canada, Winnipeg, Manitoba, Canada

David Deslauriers [1]

- Fisheries and Oceans Canada, Winnipeg, Manitoba, Canada

Ken M Jeffries

- University of Manitoba, Winnipeg, Manitoba, Canada

**Corresponding Author**

Christina M Schaefer:


### Supplementary Material


[1] Université du Québec à Rimouski, Rimouski, Quebec, Canada

**Table S1.** Raw and averaged salinity values (psu) acquired from three separate studies conducted in Frobisher Bay, Nunavut, Canada between 2008-2019.

| Collection Date | Depth (m) | Salinity Values (psu) |
|---|---|---|
| July – September 2008/2009[a] | 1m (intertidal) | 12.1 (average) |
| July – September 2008/2009[a] | 1.5m (intertidal) | 15.5 (average) |
| July – September 2008/2009[a] | 6.5m (intertidal) | 31.9 (average) |
| July – September 2008/2009[a] | 0.5m (subtidal) | 28.7 (average) |
| July – September 2008/2009[a] | 1.5m (subtidal) | 30.1 (average) |
| July – September 2008/2009[a] | 6.5m (subtidal) | 32.1 (average) |
| July – September 2008/2009[a] | 20.0m (subtidal) | 32.2 (average) |
| June 2013[b] | 0m | 32.556 |
| June 2013[b] | 0.5m | 32.53 |
| June 2013[b] | 1m | 32.478 |
| June 2013[b] | 1.5m | 32.426 |
| June 2013[b] | 2m | 32.405 |
| June 2013[b] | 2.5m | 32.41 |
| June 2013[b] | 3m | 32.416 |
| June 2013[b] | 3.5m | 32.41 |
| July 2013[b] | 0m | 32.459 |
| July 2013[b] | 0.5m | 32.426 |
| July 2013[b] | 1m | 32.424 |
| July 2013[b] | 1.5m | 32.429 |
| July 2013[b] | 2m | 32.438 |
| July 2013[b] | 2.5m | 32.441 |
| July 2013[b] | 3m | 32.439 |
| July 2013[b] | 3.5m | 32.438 |
| September 2013[b] | 0m | 32.771 |
| September 2013[b] | 0.5m | 32.611 |
| September 2013[b] | 1m | 32.452 |
| September 2013[b] | 1.5m | 32.392 |
| September 2013[b] | 2m | 32.354 |
| September 2013[b] | 2.5m | 32.319 |
| September 2013[b] | 3m | 32.301 |
| September 2013[b] | 3.5m | 32.29 |
| November 2013[b] | 0m | 32.715 |
| November 2013[b] | 0.5m | 32.514 |
| November 2013[b] | 1m | 32.455 |
| November 2013[b] | 1.5m | 32.394 |
| November 2013[b] | 2m | 32.361 |
| November 2013[b] | 2.5m | 32.337 |
| November 2013[b] | 3m | 32.312 |
| November 2013[b] | 3.5m | 32.32 |
| August 3, 2018[c] | 1m | 29.6908 |
| August 3, 2018[c] | 1m | 30.6092 |

| | | |
|---|---|---|
| August 3, 2018[c] | 1m | 31.1887 |
| August 14, 2018[c] | 1m | 30.7216 |
| August 14, 2018[c] | 1m | 29.0919 |
| August 14, 2018[c] | 1m | 30.5155 |
| August 17, 2018[c] | 1m | 29.6586 |
| August 17, 2018[c] | 1m | 29.9986 |
| August 17, 2018[c] | 1m | 30.9904 |
| August 22, 2018[c] | 1m | 25.7011 |
| August 22, 2018[c] | 1m | 31.5870 |
| August 22, 2018[c] | 1m | 31.0744 |
| August 31, 2018[c] | 1m | 30.8989 |
| August 31, 2018[c] | 1m | 31.5836 |
| August 31, 2018[c] | 1m | 31.3659 |
| September 6, 2018[c] | 1m | 31.3700 |
| September 6, 2018[c] | 1m | 30.4450 |
| September 6, 2018[c] | 1m | 31.2534 |
| September 15, 2018[c] | 1m | 30.3414 |
| September 16, 2018[c] | 1m | 31.5767 |
| September 16, 2018[c] | 1m | 31.8694 |
| September 17, 2018[c] | 1m | 31.7344 |
| September 17, 2018[c] | 1m | 31.2730 |
| September 17, 2018[c] | 1m | 32.0461 |
| September 25, 2018[c] | 1m | 32.1634 |
| September 25, 2018[c] | 1m | 32.0037 |
| September 25, 2018[c] | 1m | 32.0834 |
| October 2, 2018[c] | 1m | 32.2043 |
| October 2, 2018[c] | 1m | 32.1320 |
| October 2, 2018[c] | 1m | 32.1844 |
| October 2, 2018[c] | 1m | 32.2398 |
| July 19, 2019[d] | 1m | 31.2784 |
| July 19, 2019[d] | 1m | 31.1405 |
| July 30, 2019[d] | 1m | 30.6953 |
| July 30, 2019[d] | 1m | 30.5608 |
| July 30, 2019[d] | 1m | 30.9143 |
| August 6, 2019[d] | 1m | 31.3291 |
| August 6, 2019[d] | 1m | 31.4742 |
| August 6, 2019[d] | 1m | 31.2092 |
| August 12, 2019[d] | 1m | 31.5882 |
| August 12, 2019[d] | 1m | 31.1103 |
| August 12, 2019[d] | 1m | 31.0679 |
| August 19, 2019[d] | 1m | 31.4768 |
| August 19, 2019[d] | 1m | 31.1464 |
| August 19, 2019[d] | 1m | 31.2788 |
| August 26, 2019[d] | 1m | 31.7488 |
| August 26, 2019[d] | 1m | 31.7502 |
| August 26, 2019[d] | 1m | 31.8020 |

| | | |
|---|---|---|
| September 2, 2019[d] | 1m | 30.5557 |
| September 2, 2019[d] | 1m | 31.6892 |
| September 2, 2019[d] | 1m | 31.5706 |
| September 9, 2019[d] | 1m | 32.1259 |
| September 10, 2019[d] | 1m | 31.1015 |
| September 10, 2019[d] | 1m | 32.1029 |
| September 12, 2019[d] | 1m | 32.1449 |
| September 12, 2019[d] | 1m | 31.9427 |
| September 12, 2019[d] | 1m | 31.9947 |
| September 16, 2019[d] | 1m | 31.6366 |
| September 16, 2019[d] | 1m | 28.9074 |
| September 16, 2019[d] | 1m | 30.8813 |
| September 23, 2019[d] | 1m | 32.0937 |
| September 23, 2019[d] | 1m | 31.9992 |
| September 23, 2019[d] | 1m | 29.8791 |

[a] Spares, A.D., Stokesbury, M.J.W., O'Dor, R.K., Dick, T.A., 2012. Temperature, salinity, and prey availability shape the marine migration of Arctic char, Salvelinus alpinus, in a macrotidal estuary. Mar. Biol. 159, 1633–1646. https://doi.org/10.1007/s00227-012-1949-y

[b] Bannister, C., Snarby, A., Cheater, J., Ishulutaq, L., 2013. RV Nuliajuk Seabed Mapping Cruise Report. University of New Brunswick.

[c] Fisheries and Oceans Canada, 2018. Coastal Environmental Baseline Project (Salinity). Department of Fisheries and Oceans, Frobisher Bay, Nunavut.

[d] Fisheries and Oceans Canada, 2019. Coastal Environmental Baseline Project (Salinity). Department of Fisheries and Oceans, Frobisher Bay, Nunavut.

**Table S2.** Ranges of LA-ICP-MS operating conditions and data-acquisition parameters.

| | |
|---|---|
| Laser Ablation | |
| Model | New Wave UP-213 |
| Beam diameter (µm) | 55 |
| Energy density on sample (J $cm^{-2}$) | ~6 |
| Repetition rate (Hz) | 20 |
| Preablation time (s) | 30 |
| Ablation speed (µm $s^{-1}$) | 10 |
| ICP - MS | |
| Model | Thermo-Finnigan Element 2 |
| Sample time (ms) | 5 |
| Plasma power (W) | 1265 |
| Argon flow rate | |
| carrier gas (He) (L $min^{-1}$ ) | 0.86 |
| auxiliary gas (Ar) (L $min^{-1}$ ) | 0.78 |
| sample (L $min^{-1}$ ) | 0.83 |
| Dwell time (s) | 30 |
| Samples per peak | 5 |
| Integration window | 60% |
| Scan method | E-Scan |

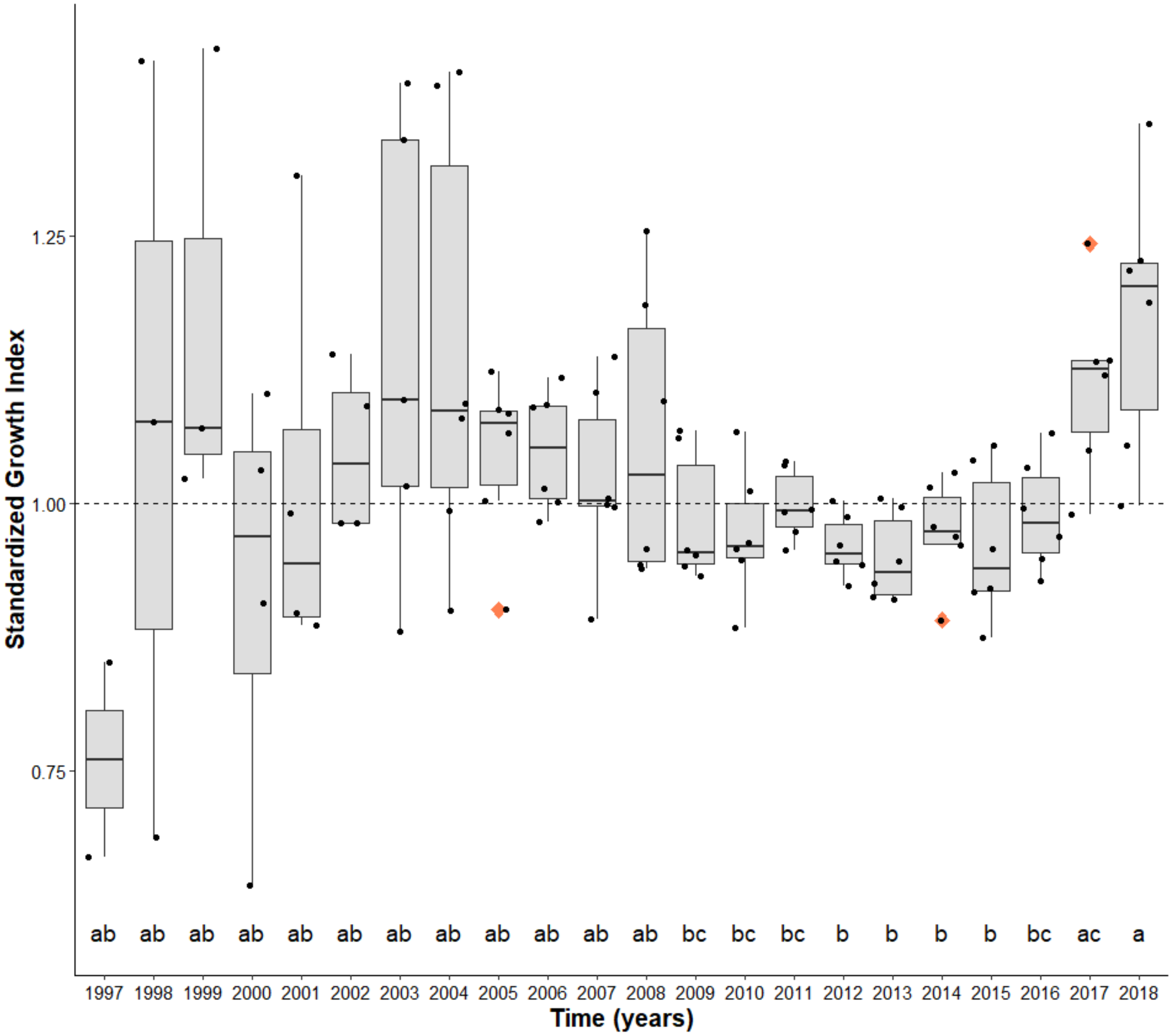


**Figure S1.** Standardized growth index (± s.e.m.) for *Mya truncata* from all sampling locations within Inner Frobisher Bay, Nunavut, Canada indicating variation in annual growth unrelated to age. The y-axis is unitless and the dashed line in each plot represents an SGI of 1.0 with values above indicating better than expected years of growth and values below the line representing less than expected growth for those years. Lowercase letters denote significant temporal differences using LMM and Tukey's HSD post-hoc test (α < 0.05). Coral colored diamonds represent outliers.

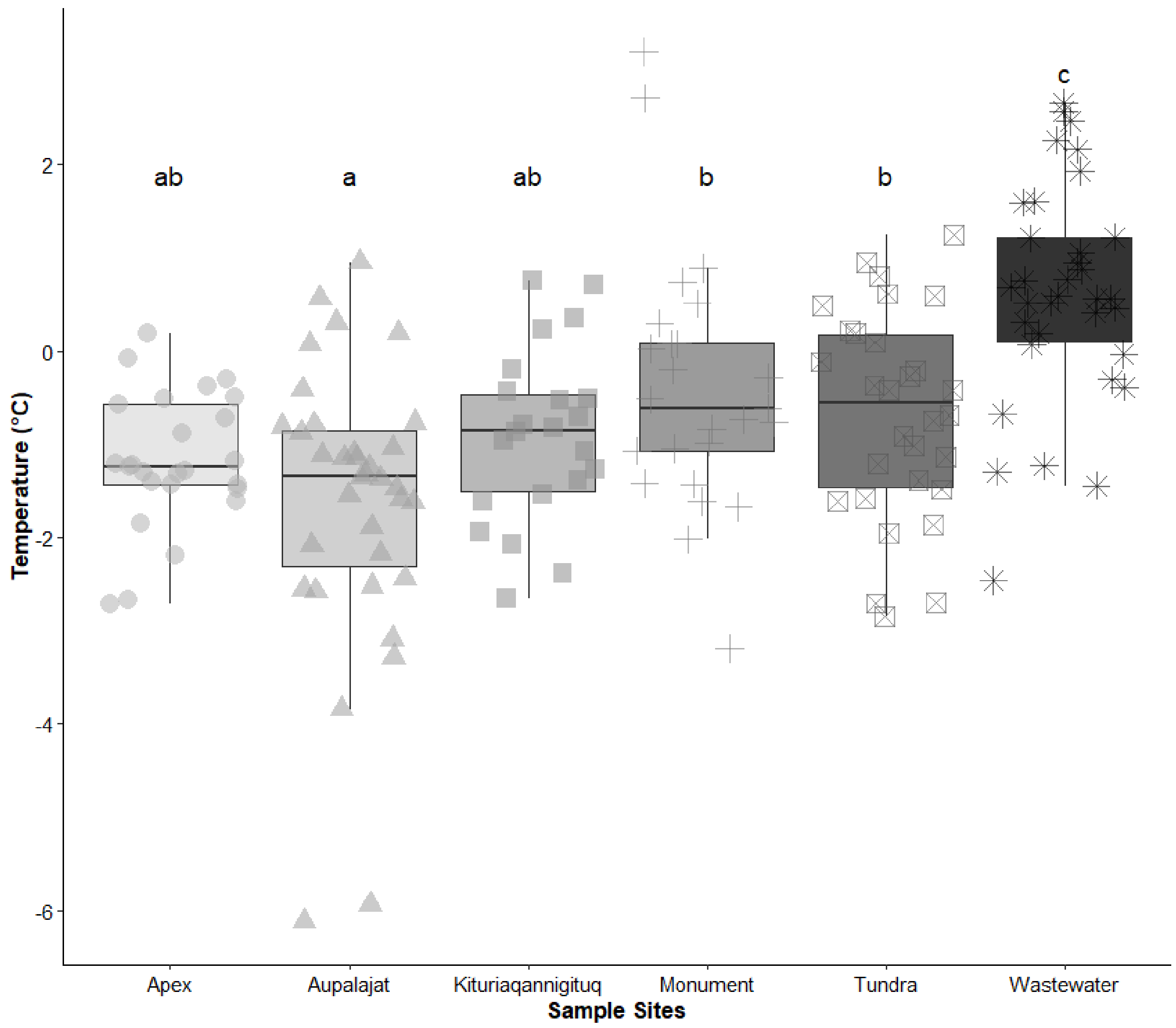


**Figure S2.** Reconstructed sea surface temperatures (°C) from *Mya truncata* shell $\delta^{18}O$ (‰ VPDB) and seawater $\delta^{18}O$ (‰ VSMOW) concentrations. Black error bars represent ± s.e.m. and all samples were collected within Inner Frobisher Bay, Nunavut, Canada. Lower case letters denote significant differences in temperature between locations using one-way ANOVA and Tukey's HSD post-hoc test ($\alpha < 0.05$).

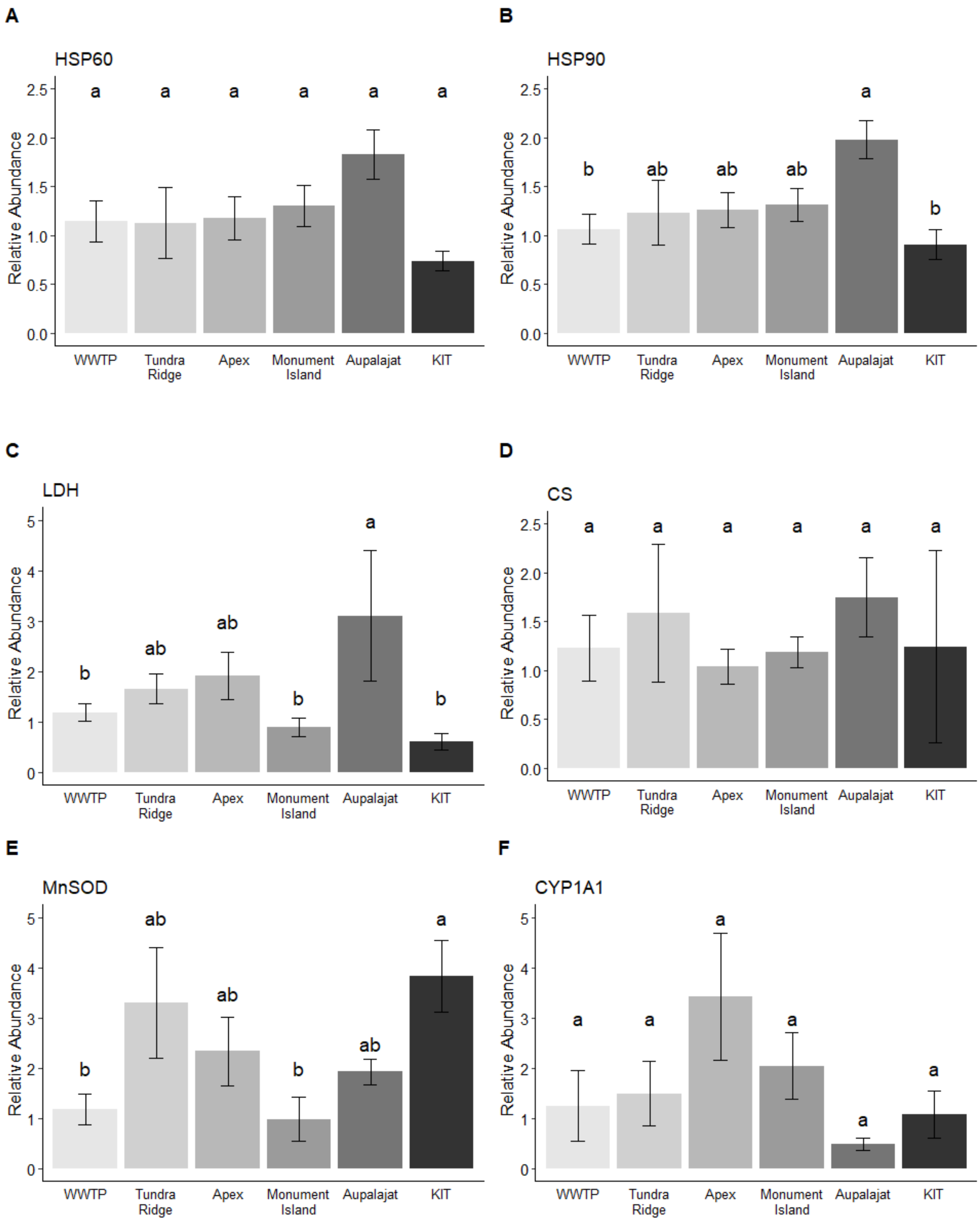
A
HSP60
B
HSP90
C
LDH
D
CS
E
MnSOD
F
CYP1A1
Relative Abundance
WWTP
Tundra Ridge
Apex
Monument Island
Aupalajat
KIT

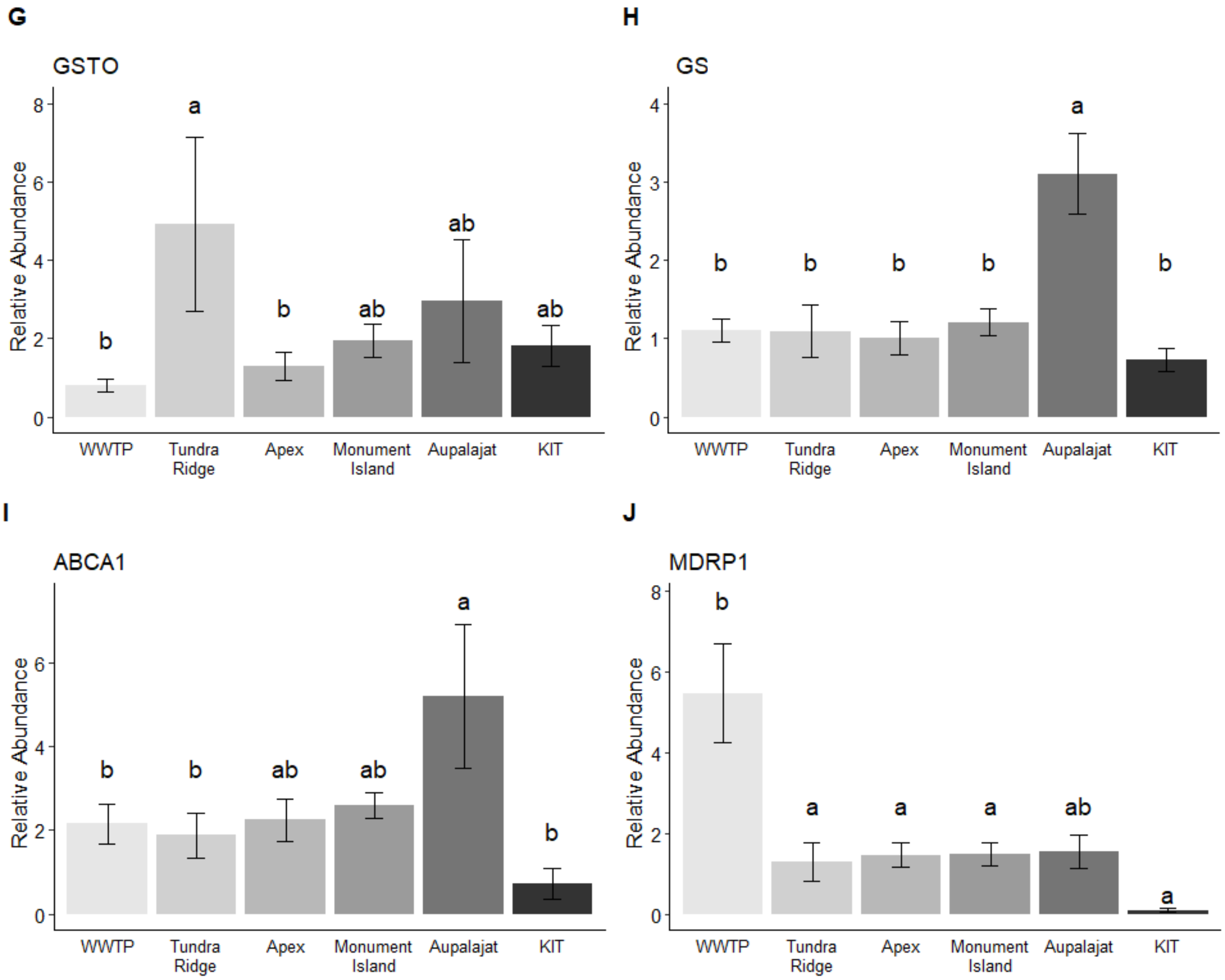


**Figure S3.** Relative expression of mRNA in the gill tissue of *Mya truncata* (± s.e.m.) organisms for 10 genes. All samples were collected in Inner Frobisher Bay, Nunavut, Canada. Lowercase letters denote statistical significance between sampling locations as determined by a one-way ANOVA and Tukey's HSD post-hoc test ($\alpha < 0.05$). KIT represents Kituriaqannigituq and WWTP represents the wastewater treatment plant.

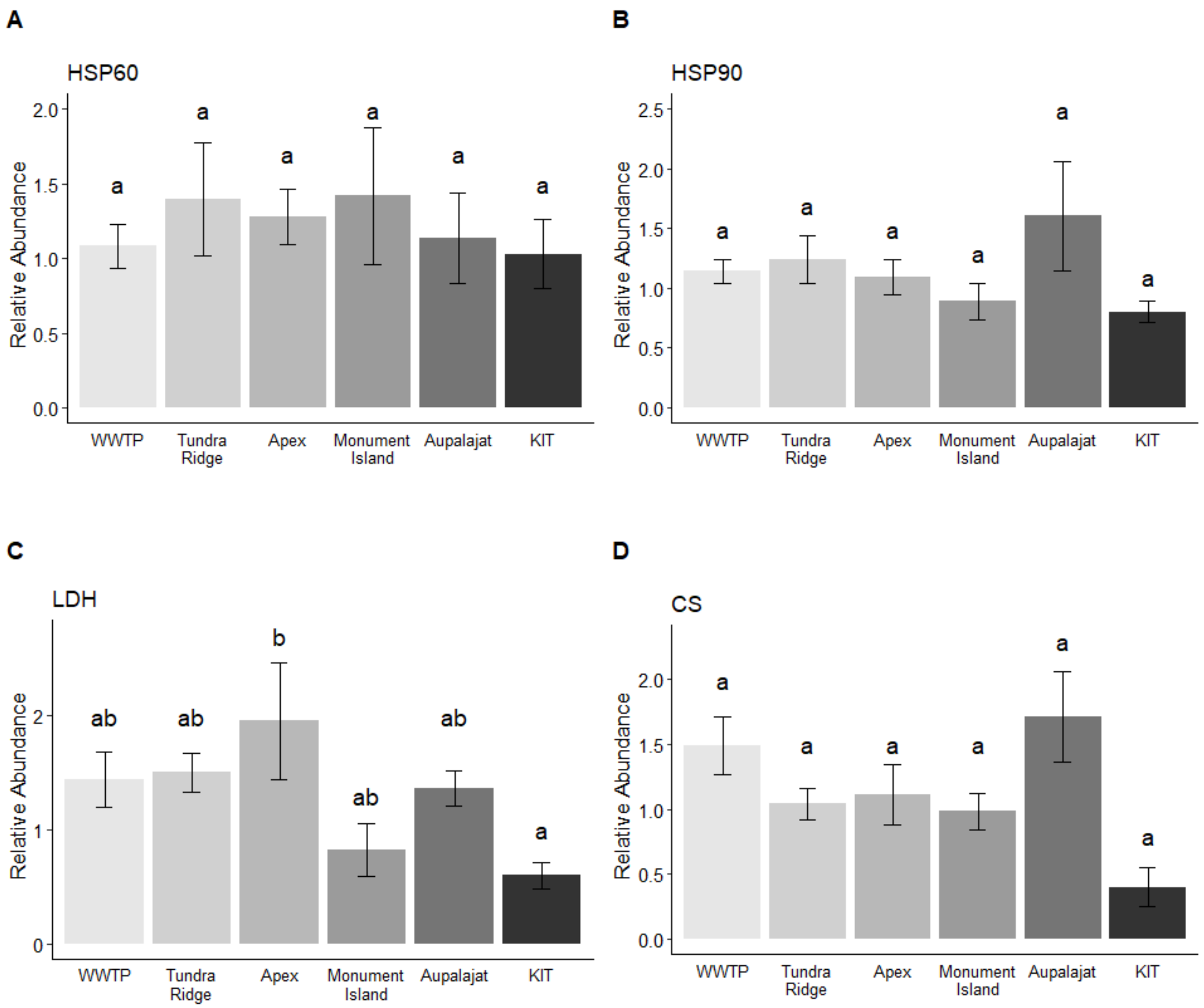
A
HSP60
Relative Abundance
2.0
1.5
1.0
0.5
0.0
a
a
a
a
a
a
WWTP
Tundra Ridge
Apex
Monument Island
Aupalajat
KIT
B
HSP90
Relative Abundance
2.5
2.0
1.5
1.0
0.5
0.0
a
a
a
a
a
a
WWTP
Tundra Ridge
Apex
Monument Island
Aupalajat
KIT
C
LDH
Relative Abundance
2
1
0
ab
ab
b
ab
ab
a
WWTP
Tundra Ridge
Apex
Monument Island
Aupalajat
KIT
D
CS
Relative Abundance
2.0
1.5
1.0
0.5
0.0
a
a
a
a
a
a
WWTP
Tundra Ridge
Apex
Monument Island
Aupalajat
KIT

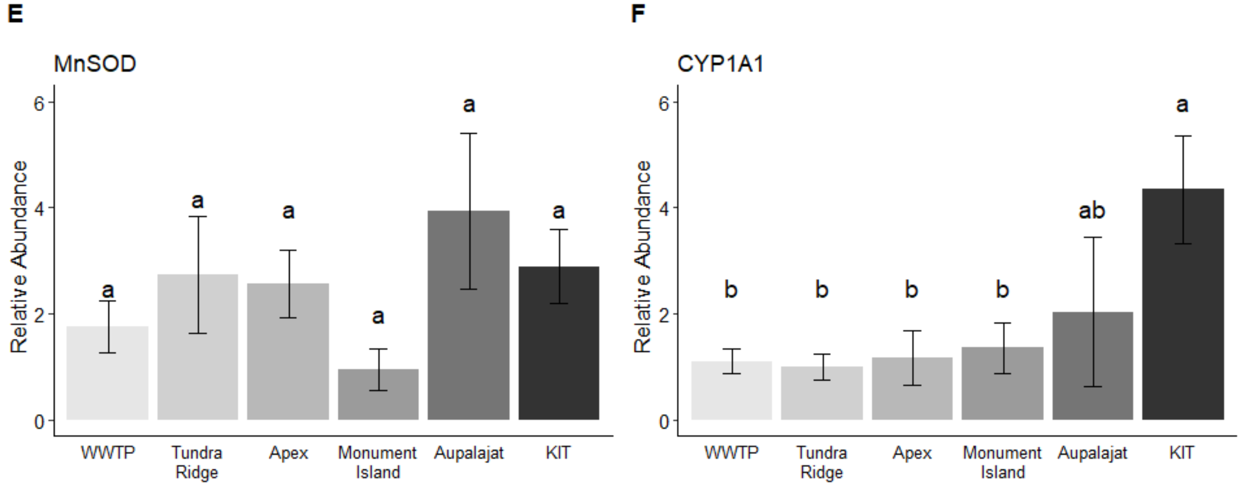
E
MnSOD
Relative Abundance
6
4
2
0
a
a
a
a
a
a
WWTP
Tundra Ridge
Apex
Monument Island
Aupalajat
KIT
F
CYP1A1
Relative Abundance
6
4
2
0
b
b
b
b
ab
a
WWTP
Tundra Ridge
Apex
Monument Island
Aupalajat
KIT

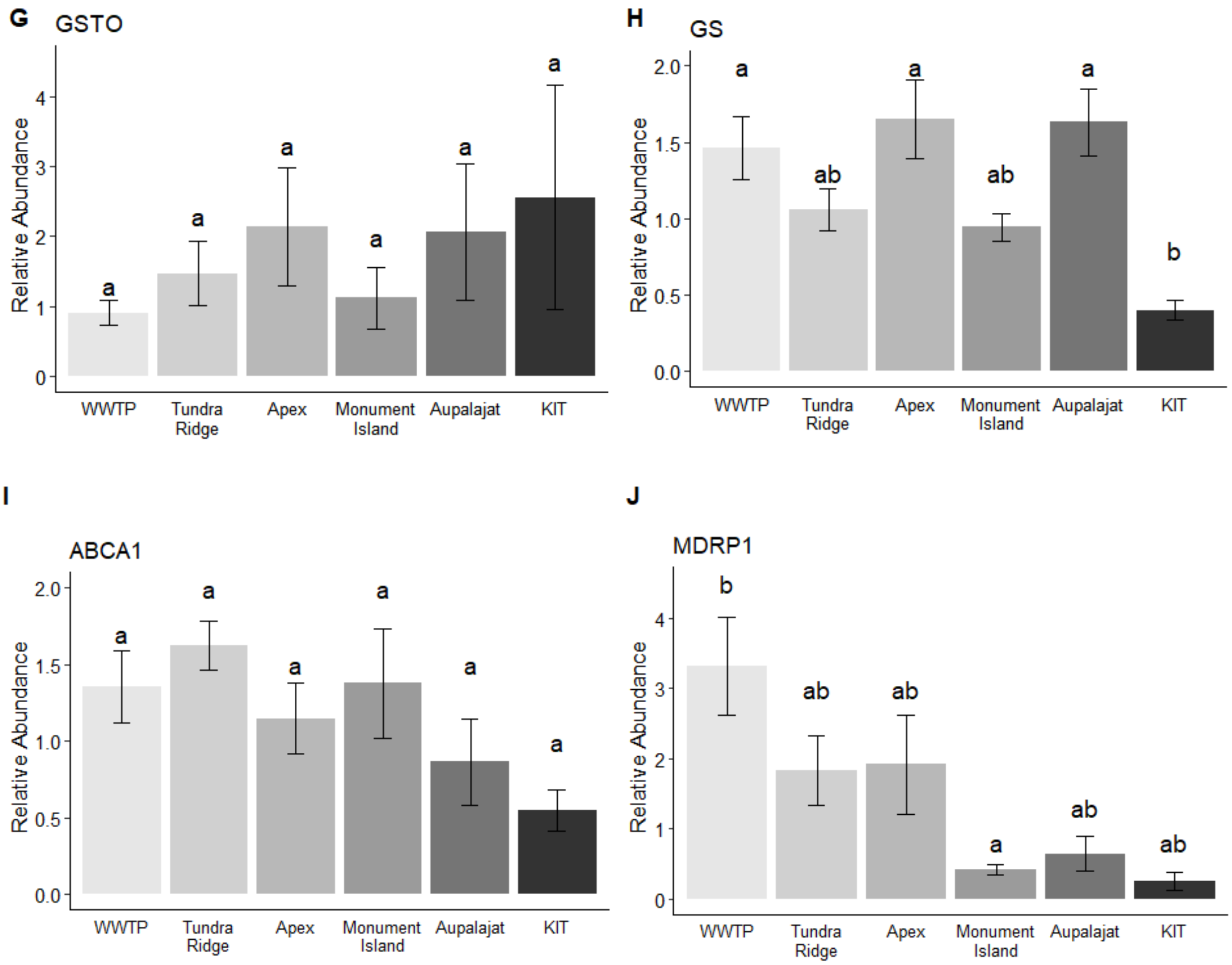


**Figure S4.** Relative expression of mRNA in the mantle tissue of *Mya truncata* (± s.e.m.) organisms for 10 genes. All individuals were collected within Inner Frobisher Bay, Nunavut, Canada. Lowercase letters denote statistical significance between sampling sites as determined by a one-way ANOVA and Tukey's HSD post-hoc test ($\alpha < 0.05$). KIT represents Kituriaqannigituq and WWTP represents the wastewater treatment plant.
